# A size principle for leg motor control in *Drosophila*

**DOI:** 10.1101/730218

**Authors:** Anthony W Azevedo, Evyn S Dickinson, Pralaksha Gurung, Lalanti Venkatasubramanian, Richard Mann, John C Tuthill

## Abstract

To move the body, the brain must precisely coordinate patterns of activity among diverse populations of motor neurons. In many species, including vertebrates, the motor neurons innervating a given muscle fire in a specific order that is determined by a gradient of cellular size and electrical excitability. This hierarchy allows premotor circuits to recruit motor neurons of increasing force capacity in a task-dependent manner. However, it remains unclear whether such a size principle also applies to species with more compact motor systems, such as the fruit fly, *Drosophila melanogaster*, which has just 53 motor neurons per leg. Using *in vivo* calcium imaging and electrophysiology, we found that genetically-identified motor neurons controlling flexion of the fly tibia exhibit a gradient of anatomical, physiological, and functional properties consistent with the size principle. Large, fast motor neurons control high force, ballistic movements while small, slow motor neurons control low force, postural movements. Intermediate neurons fall between these two extremes. In behaving flies, motor neurons are recruited in order from slow to fast. This hierarchical organization suggests that slow and fast motor neurons control distinct motor regimes. Indeed, we find that optogenetic manipulation of each motor neuron type has distinct effects on the behavior of walking flies.

## Introduction

Dexterous motor behaviors require precise neural control of muscle contraction to coordinate force production and timing across dozens to hundreds of muscles. This coordination is mediated by populations of motor neurons, which translate commands from the central nervous system into dynamic patterns of muscle contraction. Although motor neurons are the final common output of the brain, the scale and complexity of many motor systems have made it challenging to understand how motor neuron populations collectively control muscles and thus generate behavior. For example, a human leg is innervated by over 150,000 motor neurons and a single calf muscle is innervated by over 600 motor neurons (Kernell, 2006). How can the nervous system coordinate the activity of such large motor neuron populations while maintaining speed, flexibility, and precision? Is the activity of each motor neuron independently specified, or do there exist organizational principles that help reduce the dimensionality of neuromuscular control?

One way to streamline motor control is to establish a hierarchy among neurons controlling a particular movement, such as flexion of a joint. This hierarchy allows premotor circuits to excite different numbers of motor neurons depending on the required force: motor neurons controlling slow or weak movements are recruited first, followed by motor neurons that control progressively stronger, faster movements (Denny-Brown, 1929). A recruitment order for vertebrate motor neurons innervating a single muscle was first postulated over 60 years ago (Henneman, 1957). Subsequent work identified mechanisms associated with the recruitment order and synthesized these findings as the *size principle*, which states that small motor neurons, with lower spike thresholds, are recruited prior to larger neurons, which have higher spike thresholds (Henneman and Olson, 1965; Henneman et al., 1965a; Henneman et al., 1965b; Mcphedran et al., 1965; Mendell, 2005; Wuerker et al., 1965). Evidence for the size principle has been provided by electrophysiological analysis of motor neurons in a number of species, including humans (Milner-Brown et al., 1973). These studies have also described systematic relationships between motor neuron electrical excitability, recruitment order, conduction velocity, force production, and strength of sensory feedback and descending input (Bawa et al., 1984; Binder et al., 1983; Fleshman et al., 1981; Kernell and Sjöholm, 1975; Zengel et al., 1985). Although there exist notable exceptions (Desmedt and Godaux, 1981; Menelaou and McLean, 2012; Smith et al., 1980), the size principle and hierarchical recruitment order nonetheless provide a useful framework for understanding how premotor circuits efficiently coordinate large pools of motor neurons.

The leg of the fruit fly, *Drosophila melanogaster*, contains 14 muscles which are innervated by just 53 motor neurons (Baek and Mann, 2009; Brierley et al., 2012; Soler, 2004; Maniates-Selvin et al., 2019). This small scale raises the possibility that flies may require a distinct strategy for limb motor control that does not rely on a strict recruitment hierarchy (Belanger, 2005). Previous studies in the locust and crayfish found gradients in motor neuron properties that resemble the size principle (Gabriel et al., 2003; Hill and Cattaert, 2008; Sasaki and Burrows, 1998). However, these species also possess inhibitory GABAergic motor neurons, which are thought to extend the level of control over leg muscle dynamics (Wolf, 2014). *Drosophila* lack inhibitory motor neurons (Schmid et al., 1999; Witten and Truman, 1998), raising the question of whether the organization of the fly leg motor system is fundamentally different.

Understanding motor control in *Drosophila* is important because the fly’s compact nervous system and identified cell types make it a tractable system for comprehensive circuit analysis. However, up to this point, the functional architecture of the *Drosophila* leg motor control system has not been studied. While a great deal of progress has been made on understanding the processing of sensory signals in the *Drosophila* brain, comparatively little is known about how this information is translated into behavior by motor circuits in the fly’s ventral nerve cord (VNC).

Here, we use *in vivo* calcium imaging, whole-cell patch clamp electrophysiology, and optogenetics during walking behavior to investigate motor control of the *Drosophila* tibia (**Figure 1A**). The femur-tibia joint of a walking fly flexes and extends 10-20 times per second, reaching swing speeds of several thousand degrees per second (DeAngelis et al., 2019; Gowda et al., 2018; Mendes et al., 2013; Strauss and Heisenberg, 1990; Wosnitza et al., 2013). Flies also use their legs to target other body parts during grooming (Hampel et al., 2015; Seeds et al., 2014). Thus, motor control of the tibia must be both fast and precise.

**Figure 1.**
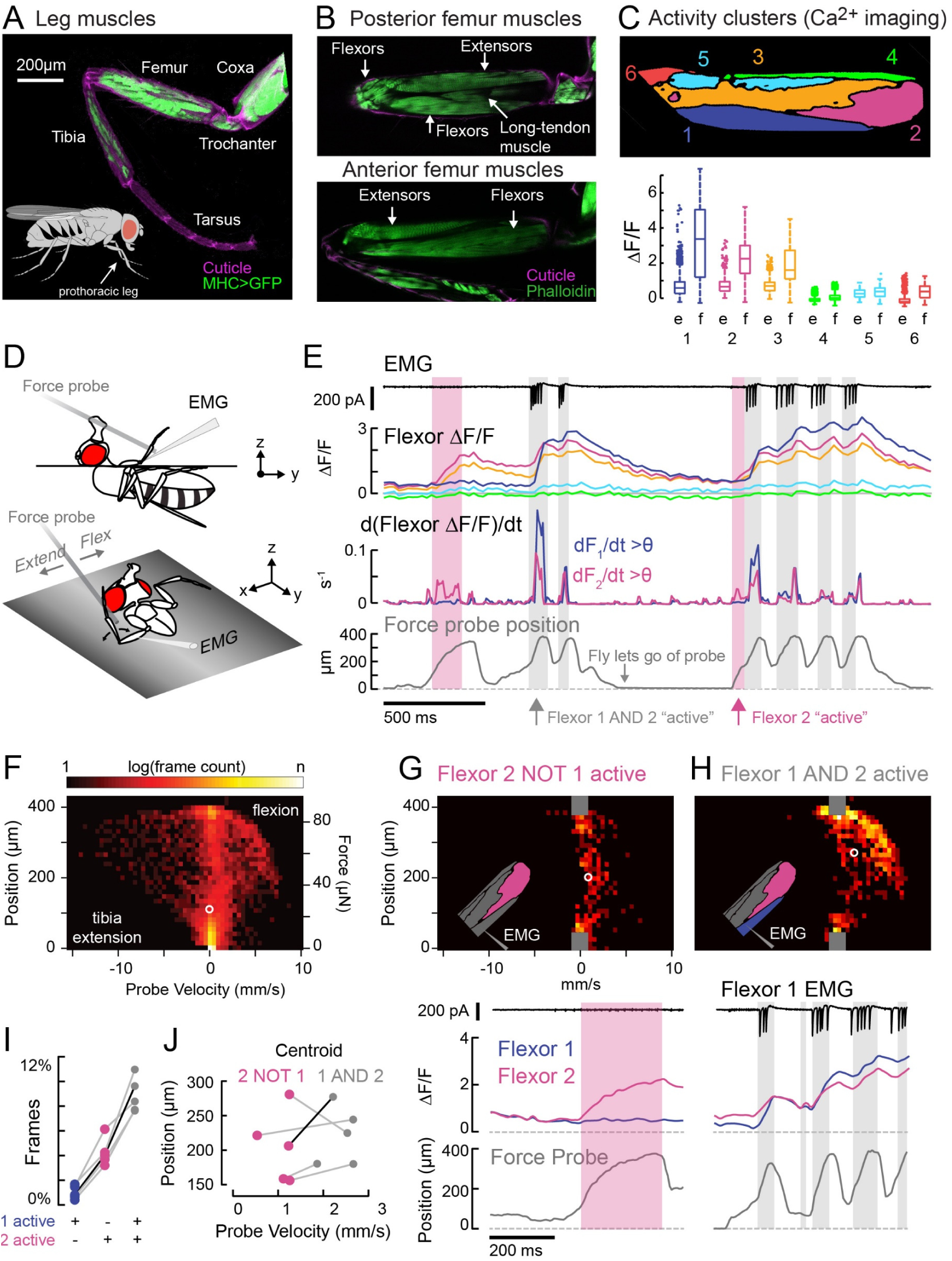
Functional organization and recruitment order among muscles controlling the *Drosophila* leg. **A.** Muscles of the right prothoracic leg of a female *Drosophila* (*MHC-LexA; 20XLexAop-GFP*). **B.** Muscles controlling tibia movement in the fly femur. Top and bottom are confocal sections through the femur. Anterior-posterior axis refers to a leg positioned to the side of the body (Soler et al., 2004). **C.** Top: K-means clustering of calcium signals (*MHC-GAL4;UAS-Gcamp6f*) based on correlation of pixel intensities across 180 s of leg movements in an example fly. Bottom: average change in fluorescence for each cluster, in each frame, when the leg is extended (femur-tibia joint >120°, 20772 frames) vs. flexed (<30°, 61,860 frames), n = 5 flies. Responses to flexion were consistently higher (p=0.01 for cluster 5, p<10^-6^ for all other clusters, 2-way ANOVA, Tukey-Kramer correction). **D.** Schematic of the experimental setup. The fly is fixed in a holder so that it can pull on a calibrated force probe with the tibia while calcium signals are recorded from muscles in the femur. **E.** Calcium activity in tibia muscles while the fly pulls on the force probe (bottom trace). Cluster 6 was obscured by the probe and not included. The middle row shows the smoothed, rectified derivative of the cluster fluorescence (dF_i_/dt) for the two brightest clusters (1 & 2), which we refer to as Flexors. Highlighted periods indicate that both Flexors 1 and 2 are active simultaneously (gray, dF/dt > 0.005), or that Flexor 2 alone is active (magenta). **F.** 2D histogram of probe position and velocity, for all frames (n = 13,504) for a representative fly. The probe was often stationary (velocity=0), either because the fly let go of the probe (F = 0), or because the fly pulled the probe as far as it could (F ∼85 µN), reflected by the hotspots in the 2D histogram. In F-H, the white circles indicate the centroids of the distributions. **G.** 2D histogram of probe position vs. velocity when Flexor 2 fluorescence increased, but not Flexor 1 (n = 637 frames, same fly as F). Gray squares indicate hotspots in F, which are excluded here. Color scaled to log(50 frames). **H.** Same as G, when Flexor 1 AND 2 fluorescence increased simultaneously (n = 1,449 frames). **I.** Fraction of total frames for each fly in which both Flexor 1 and 2 fluorescence increased (gray), Flexor 2 alone increased (magenta), or Flexor 1 alone increased. Number of frames for each of five flies: 32,916, 13,504, 37,136, 37,136, 24,476. **J.** Shift in the centroid of the 2D histogram when Flexor 1 fluorescence is increasing along with Flexor 2 fluorescence (gray), compared to when Flexor 2 fluorescence alone is increasing (p<0.01, Wilcoxon rank sum test). Black line indicates example cell in G and H.

We found that motor neurons that control tibia flexion lie along a gradient of anatomical, functional, and intrinsic electrophysiological properties. *Slow* motor neurons have thin axons, low spike thresholds, and produce little force per spike, while *fast* motor neurons have wider axons, higher spike thresholds, and produce large forces with just a single spike. During spontaneous leg movements, tibia motor neurons follow a recruitment hierarchy: slow motor neurons are recruited first, followed by intermediate, then fast neurons. These results are consistent with the size principle, suggesting that leg motor control in the fly is organized hierarchically, similar to other limbed animals, including vertebrates.

## Results

### Organization and recruitment of motor units controlling the femur-tibia joint of Drosophila

We first sought to understand the relationship between muscle activity and movement of the fly tibia (**Figure 1B**). Fly leg muscles are each composed of multiple fibers (Soler, 2004) and innervated by distinct motor neurons (Baek and Mann, 2009; Brierley et al., 2012). A motor neuron and the muscle fibers it innervates are referred to as a *motor unit*. In most invertebrate species, multiple motor neurons can innervate the same muscle fiber, so motor units may be overlapping (Hoyle, 1983).

We used wide-field fluorescence imaging of muscle calcium signals with GCamp6f (Chen et al., 2013; Lindsay et al., 2017) to map the spatial organization of motor units in the fly femur during spontaneous tibia movements (**Figures 1C and S1; Movie S1**). We observed that spatial patterns of calcium activity were different for different movements (e.g., tibia flexion vs. extension).

To quantify this spatial organization, we performed unsupervised clustering so that pixels with correlated activity were grouped together into *activity clusters*. Three clusters significantly increased their fluorescence when the leg was flexed (**Figure 1C**); we refer to these clusters as Flexors 1, 2, and 3. Flexors 2 and 3 were active during most periods of tibia flexion, whereas Flexor 1 was only active only during large, fast movements (**Figure 1E**). This organization was similar across flies (**Figure S1C**) and robust to changes in clustering parameters (**Figure S1F**). Likely because of the orientation of the leg in these experiments, we did not observe any clusters whose activity increased specifically during tibia extension (see **Figure S1D-E** and Experimental Procedures for further discussion). For the remainder of the study, we focus our efforts on motor control of tibia flexion.

For Flexors 1 and 2, we could record and identify extracellular motor neuron spikes in the femur (electromyogram: EMG), which confirmed that the activity clusters identified through calcium imaging correspond to distinct motor units (**Figure 1E** and see **Figure 2D**, below). In other words, we propose that activity clusters 1 and 2 reflect the firing patterns of specific motor neurons.

**Figure 2.**
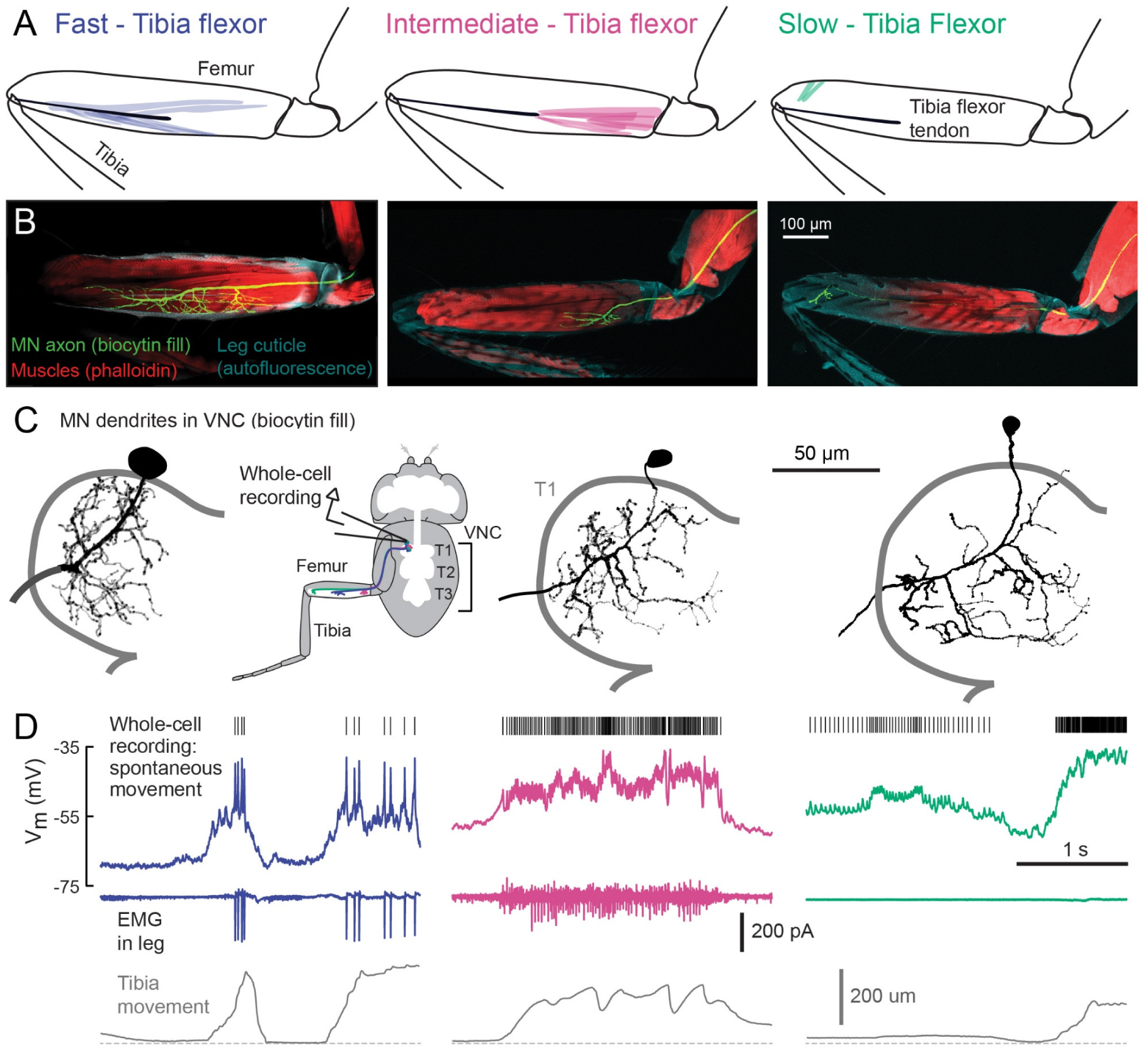
Motor neurons controlling tibia flexion. **A.** Schematic of the muscle fibers innervated by motor neurons labeled by the following Gal4 lines: left: *R81A07-Gal4*, center: *R22A08-Gal4*, and right: *R35C09-Gal4*. **B.** Biocytin fills of leg motor neurons from whole-cell patch clamp recordings, labeled in fixed tissue with strepavidin-Alexa-568 (green), counter stained with phalloidin-Alexa-633 (red) labeling actin in muscles, and autofluorescence of the cuticle (cyan). **C.** Images of biocytin fills in the VNC. Prothoracic neuromere is outlined. Cells were traced and “filled” with the simple neurite tracer plugin in Fiji. **D.** Example recordings from each motor neuron type during spontaneous tibia movement: whole-cell current clamp recordings with detected spike times above (top row, corrected for a liquid junction potential of −13 mV), EMG recordings (middle), and tibia movement (force probe position; bottom).

Muscles controlling the same joint may exert different levels of force, depending on fiber type composition and musculoskeletal biomechanics. To understand the relationship between motor unit activity and force production, we allowed the fly to pull on a flexible probe with its tibia (**Figure 1D**). We calibrated the force required to deflect the probe a given distance and measured a spring constant of 0.22 μN/μm (**Figure S2A-D**). Flies were able to move the force probe up to 400 μm, reaching speeds of 8 mm/s (**Figure 1F**). That is, the fly was capable of producing close to 100 μN of force at the tip of the tibia and changes in force of ∼1.3 mN/s. For comparison, the mass of the fly is ∼1 mg for a weight of ∼10 μN; this means that the femur-tibia joint can produce enough force to lift approximately ten times the fly’s body weight.

We used the activity clusters computed from unloaded leg movements (**Figure 1C**) to examine whether different flexor motor units control different levels of tibia force production. We observed that Flexor 2 activity increased across a range of tibia velocities and forces, but Flexor 1 activity increased only during the fastest, most powerful movements. To quantify this relationship, we compared probe force and velocity when clusters were recruited either alone or together (**Figure 1G-J**). The kinetics of GCaMP6f are slow relative to fly tibia movements, so we examined only periods when the derivative of the fluorescence signal was positive, i.e. muscle contraction was increasing (**Figure 1E**). The highest probe forces and velocities were achieved only when both Flexors 1 and 2 were active together (**Figure 1G-H**). When Flexor 2 was active alone, probe velocities were always lower (**Figure 1J**). Occasionally, the derivative of Flexor 1 fluorescence alone was high (**Figure 1I**), but in these rare instances the intensity of Flexor 2 was also high (**Figure S2E-G**), indicating that Flexor 2 was contracting. These results indicate that distinct motor units control distinct levels of force production, and that they are recruited in a specific order, with motor units controlling low forces firing prior to motor units controlling higher forces.

The results of **Figure 1** suggest two organizational features of fly leg motor control. First, the fly tibia is controlled by a number of distinct motor units that are active at different levels of force production. Second, the sequential activity of tibia flexor motor units is consistent with a hierarchical recruitment order. The spatial organization of tibia flexor motor units also provides a template that can be used to identify genetic driver lines that label specific motor neurons.

### A gradient of electrophysiological and anatomical properties among motor neurons controlling flexion of the femur-tibia joint

We visually screened a large collection of Gal4 driver lines (Jenett et al., 2012) for expression in motor neurons that innervate the activity clusters we identified through calcium imaging (**Figure 1**). We focus our analysis on three lines that each label a single tibia flexor motor neuron in the femur. The first line (*R81A07-Gal4*) labels a motor neuron that innervates the high-force motor unit (**Figure 2A-C**, left) that we identified through calcium imaging (Flexor 1). The second (*R22A08-Gal4*) labels a neuron that projects to the proximal femur (**Figure 2A-C**, middle), one of several likely candidates that controls Flexor 2 (Baek and Mann, 2009). The third line (*R35C09-Gal4*) labels a single motor neuron that projects to the distal part of the femur, where the signals in our calcium imaging experiments were weak and noisy (**Figure 2A-C**, right). The muscles in this distal region have been previously referred to as “reductors” (Baek and Mann, 2009; Brierley et al., 2012; Soler, 2004), but their alignment and attachment points suggest they control flexion of the tibia (see Experimental Procedures for further details). For consistency with the literature on other insects (Burrows, 1996), and based on their functional properties described below, we refer to these motor neurons as fast, intermediate, and slow.

The three identified motor neurons exhibit conspicuous differences in the size of their axons (**Figure 2B**), dendrites, and cell bodies (**Figure 2C, Figure S3**, and **Table S1**). The fast motor neuron has an exceptionally large soma (for a *Drosophila* neuron) and thick dendritic branches. Its axon has a wide diameter and branches extensively in the femur. The intermediate motor neuron has a smaller cell body, thinner dendritic branches, and smaller axonal arborization in the femur. The slow motor neuron has the smallest cell body, dendrites, and axon of the three. These anatomical features, measured using light microscopy, are consistent with the morphology of the same motor neurons identified in an electron microscopy volume of the fly VNC (Maniates-Selvin et al, 2019).

Using *in vivo* whole-cell patch-clamp electrophysiology, we recorded the membrane potential of each motor neuron type while the fly pulled on the force probe. The recording configuration was similar to that used for calcium imaging (**Figure 1**). We observed increases in the firing rate of each motor neuron type during tibia flexion (**Figure 2D**). Simultaneous EMG recordings from motor neuron axons in the femur confirmed that the fast and intermediate motor neurons correspond to Flexors 1 and 2 identified via calcium imaging (**Figure 1**). Perhaps because of the small size of its axon, we were unable to identify slow motor neuron spikes with extracellular recordings.

The intrinsic electrical properties of these three motor neurons varied along a continuum (**Figure 3**). The average resting potential of fast motor neurons was lower (−68 mV) than that of intermediate (−60 mV) and slow (−48 mV) motor neurons (**Figure 3B**). While the fast and intermediate neurons projecting to the flexor muscle were silent at rest, the slow neuron had a resting spike rate of approximately 30 Hz (**Figure 3C**). We also observed a gradient in input resistance: 150 MΩ for fast motor neurons, 300 MΩ for intermediate motor neurons, and 700 MΩ for slow motor neurons. Current injection in the fast and intermediate neurons failed to reliably trigger action potentials, based on recordings from both the cell body (whole-cell) and axon (EMG). This is likely because of intrinsic morphological or electrophysiological properties that electrically isolate the soma from the spike initiation zone (Sasaki and Burrows, 1998). By contrast, injecting current into the slow neuron effectively modulated the spike rate (**Figure 3A**).

**Figure 3.**
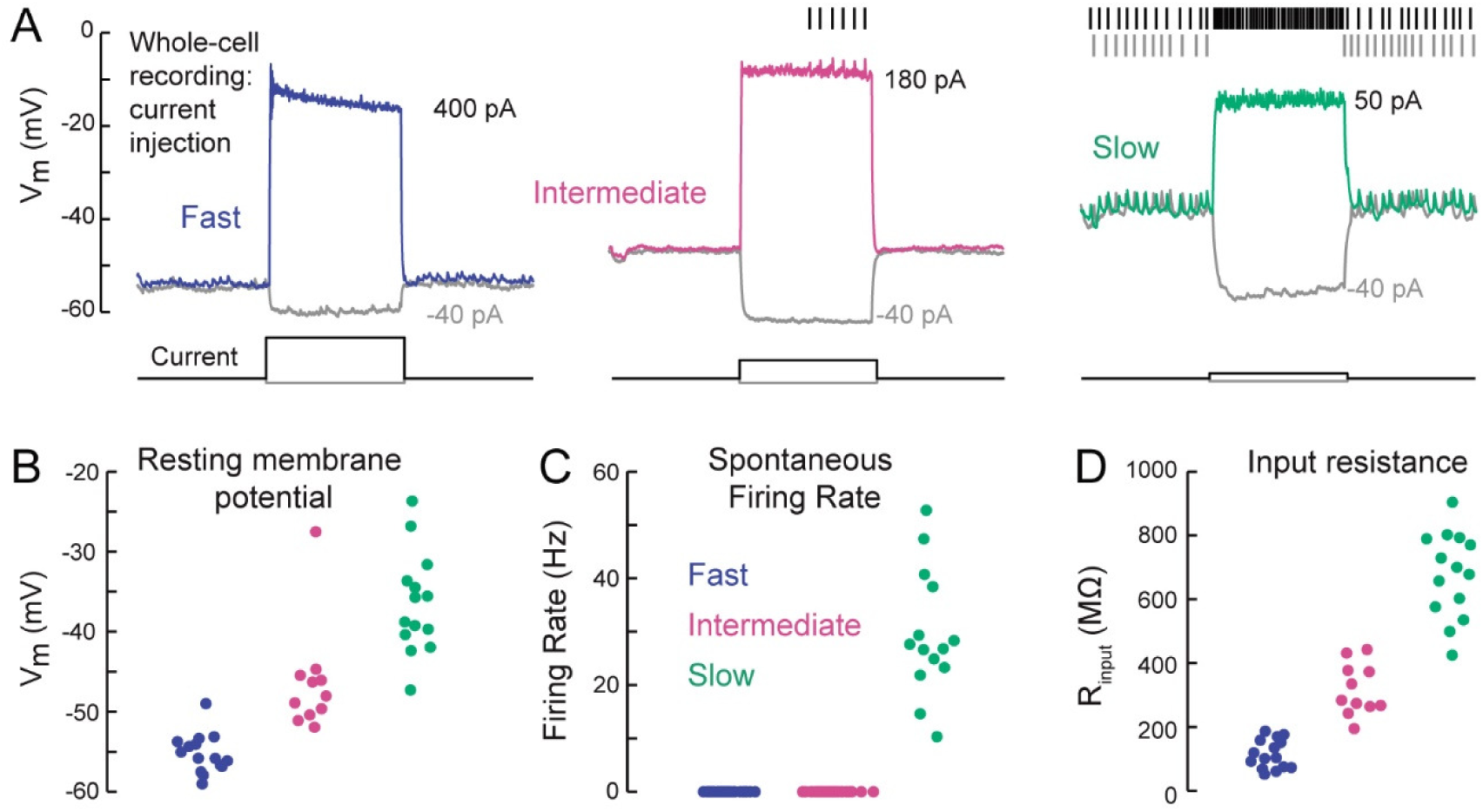
A gradient of intrinsic properties among tibia flexor motor neurons. **A.** Example traces showing voltage responses to current injection in whole-cell recordings from each motor neuron type. **B.** Average resting membrane potential for fast (n = 15), intermediate (n = 11), and slow tibia flexor motor neurons (n = 14) (p<0.001, 2-way ANOVA, Tukey-Kramer correction for multiple comparisons). **C.** Average spontaneous firing rate in fast (n = 15), intermediate (n = 11), and slow neurons (n = 14) (p<10^-5^). **D.** Input resistance in fast (n = 15), intermediate (n = 11), and slow neurons (n = 14) (p<0.001, 2-way ANOVA, Tukey-Kramer correction for multiple comparisons). All intrinsic properties in B-D were calculated from periods when the fly was not moving.

Overall, our electrophysiological characterization of tibia flexor motor neurons (**Figures 2 and 3**) reveal a systematic relationship between motor neuron anatomy, input resistance, resting membrane potential, and spontaneous firing rate. These properties accompany differences in the force generated by a spike in each neuron, which we quantify next.

### A gradient of force per spike across motor neurons

To characterize the gain, or force per spike, of each motor neuron, we measured probe displacement as a function of motor neuron firing rate. To evoke spikes in the fast and intermediate neurons, we optogenetically stimulated motor neuron dendrites in the VNC using expression of Chrimson (Klapoetke et al. 2014), a red-shifted channelrhodopsin (**Figure 4A-B**). Brief (10-20 ms) flashes of increasing intensity produced increasing numbers of spikes, corresponding spikes in the EMG, and movement of the tibia detected by small displacements of the force probe (**Movie S2**). Aligning probe movement to spike onset showed that a single fast motor neuron spike produced ∼10 μN of force, resulting in a 50 μm movement of the force probe (**Figure 4A**). An intermediate neuron spike produced ∼1 μN that moved the tibia 5 μm (**Figure 4B**). For comparison, during take-off, the peak force production of the fly leg is ∼100 μN (Zumstein 2004).

**Figure 4.**
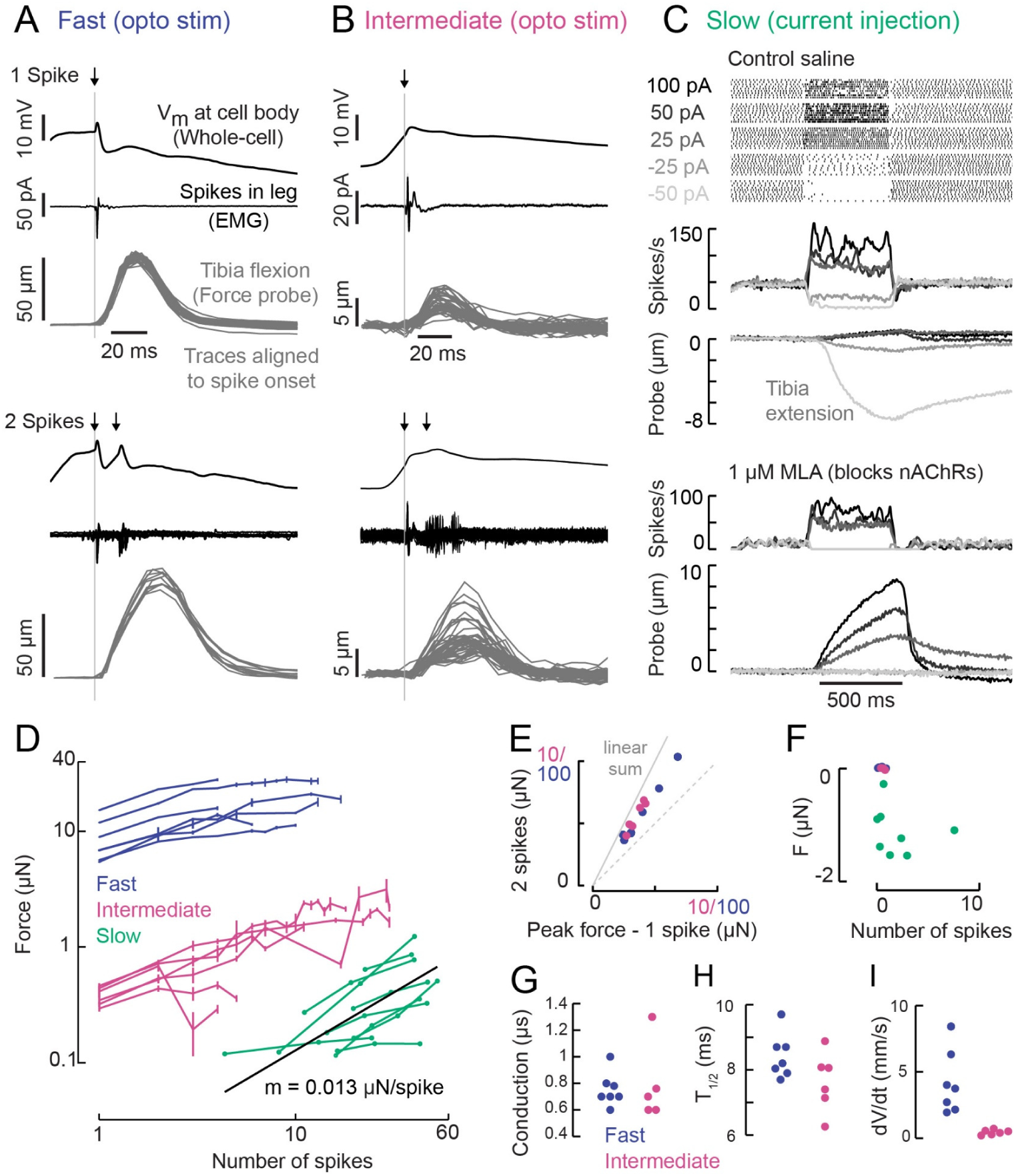
A gradient of force production among tibia flexor motor neurons. **A.** Optogenetic activation of a fast flexor motor neuron expressing Chrimson (50 ms flash from a 625 nm LED, ∼2 mW/mm^2^). Traces show average membrane potential for trials with one (top) and two (bottom) spikes, the average response in the EMG, and resulting tibia movement for each trial (50 µm = 11 µN). The jitter in the force probe movement traces results from variability of when a spike occurs relative to the video exposure (170 fps). **B.** Same as A for an example intermediate flexor motor neuron. **C.** Tibia movement resulting from current injection in a slow flexor motor neuron. Top: Spike rasters from an example cell during current injection. Firing rates are shown below, color coded according to current injection value, followed by the baseline subtracted average movement of the probe (5 µm = 1.1 µN). Bottom: spike rates and probe movement in the presence of the cholinergic antagonist MLA (1 µM), which reduces excitatory synaptic input to the motor neuron. **D.** Peak average force vs. number of spikes for fast (blue), intermediate (magenta), and slow (green) motor neurons. The number of spikes in slow neurons is computed as the average number of spikes during positive current injection steps minus the baseline firing rate, i.e. the number of additional spikes above baseline. The black line is a linear fit to the slow motor neuron data points, with the slope indicated below. **E.** Peak probe displacement for 2 spikes vs. 1 spike in fast (blue) and intermediate motor neurons (magenta). **F.** Summary data showing that zero spikes in fast (n = 7) and intermediate neurons (n = 6) does not cause probe movement, but that hyperpolarization in slow motor neurons (n = 9) causes the fly to let go of the probe, i.e. decreases the applied force. The number of spikes (x-axis) is computed as the average number of spikes per trial during the hyperpolarization.**G.** Delay between a spike in the cell body and the EMG spike (conduction delay). Note, there may be a delay from the spike initiation zone to the cell body that is not captured (n = 5 intermediate cells), p = 0.6, Wilcoxon rank sum test. **H.** Time to half maximal probe displacement for fast (blue) and intermediate cells (magenta), p = 0.2, rank sum test. **I.** Estimates of the maximum velocity of tibia movement in each fast (blue) and intermediate motor neuron (magenta), p = 0.0012, rank sum test. A line was fit to the rising phase of probe points aligned to single spikes as in B and D.

Increasing the spike rate of the slow motor neuron with current injection produced small (∼1 µm) and slow tibia movements (**Figure 4C**). Decreasing slow motor neuron firing produced tibia extension (**Figure 4C, E**), suggesting that spontaneous firing in this cell contributes to the resting force on the probe. Bath application of 1 μM MLA, an antagonist of nicotinic acetylcholine receptors, led to a decrease in the spontaneous firing rate and reduced the resting force on the probe by ∼1.5 μN, or ∼15% of the fly’s weight (**Figure 4C**). This suggests that the spontaneous firing rate in slow motor neurons is set by excitatory synaptic input.

A comparison of force production as a function of firing rate revealed that fast, intermediate and slow motor neurons occupy distinct force production regimes (**Figure 4D**). The shape of these force production curves were also distinct. For fast and intermediate neurons, the force produced by two spikes was ∼1.6X the force produced by a single spike (**Figure 4E**) and the force-per-spike curves saturated at ∼10 spikes (**Figure 4D**). These observations suggest that the fast and intermediate muscle fibers fatigue during repeated stimulation. In contrast, the resting spike rate of slow motor neurons maintains constant force on the probe (**Figure 4C, F**). Thus, slow motor neurons may be used to maintain body posture, while intermediate and fast motor neurons are transiently recruited to execute body movements. Consistent with this hypothesis, we did not observe sustained firing in EMG recordings of fast or intermediate neurons during spontaneous leg movements (**Figure 1E**).

The rates of force generation for the fast and intermediate motor units were greater than the slow motor unit. For example, the effect of a spike in the fast and intermediate motor neurons reached half maximal force in ∼8.5 ms (**Figure 4H**), whereas the effect of increasing the spike rate in slow motor neurons was gradual and did not reach its peak within 500 ms. Hyperpolarizing slow motor neurons required ∼100 ms to reach its maximal effect, which could reflect slow release of muscle tension or could be due to activity of other motor neurons in the population that influence resting force on the probe.

Several lines of evidence suggest that the fast and slow motor neurons represent the extremes of tibia flexor motor neuron size and force production. The fast neuron labeled by *81A07-Gal4* is the largest motor neuron targeting the femur (Baek and Mann, 2009; Brierley et al., 2012), and gives rise to the largest EMG spikes we observed in the leg. The slow neuron labeled by *35C09-Gal4* targets two of the most distal muscle fibers in the femur, which consequently have the smallest mechanical advantage. We also recorded from several other motor neurons controlling tibia flexion, none of which had more extreme properties. Importantly, these cells exhibited similar relationships in morphology, intrinsic properties, and force production (**Figure S4**), which provides confidence that these correlations are not an artifact of the motor neurons we have chosen to highlight in this study.

### Motor neurons are recruited in a temporal sequence from weakest to strongest

Calcium imaging from leg muscles suggested that distinct motor units are active during distinct tibia movement regimes, and that this activity follows a recruitment order (**Figure 1**). To test these hypotheses more rigorously, we examined recordings from single cells and pairs of tibia flexor motor neurons during spontaneous leg movements (**Figure 5A-C**).

**Figure 5.**
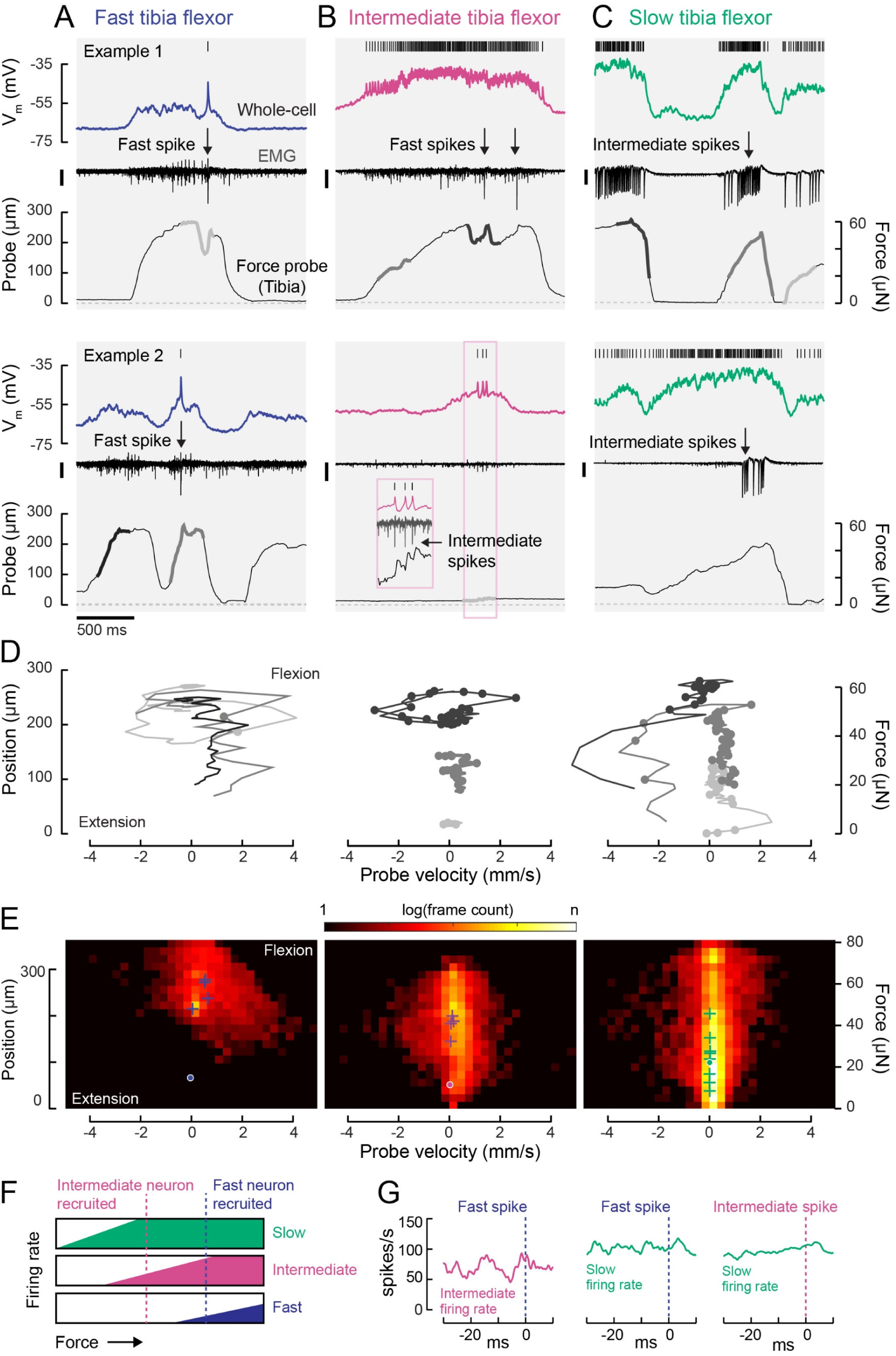
Motor neurons are recruited in a specific order across different motor regimes. **A.** Membrane potential, EMG, and probe movement, for two example epochs of spontaneous leg movement during a whole-cell recording of the fast motor neuron. Highlighted events are plotted in D. The EMG was recorded in the proximal femur to pick up both fast and intermediate spikes (scale is 10 pA). **B.** Same as A, but for an intermediate motor neuron. The EMG for the intermediate neuron was recorded in the proximal part of the femur, the largest events reflect activity in the fast motor neuron (scale is 100 pA). **C.** Same as A-B, but for a slow motor neuron. The EMG was recorded in the proximal femur and large events are likely from the intermediate neuron (scale is 100 pA). **D.** Example force probe trajectories from movement examples. The grayscale in D matches that of the trajectories in A-C. Each vertex is a single frame; vertices with dots indicate the occurrence of a spike in the whole-cell recording. **E.** 2D histograms of probe force and velocity for video frames within 25 ms following a spike across recordings in cells of each type (fast – 5 neurons, 4,666 frames; intermediate – 4 neurons, 11,833 frames; slow – 9 neurons, 65,473 frames). Crosses indicate the centroids of the histograms for individual neurons. Dots outlined in white indicate centroid of the 2D histograms of all frames. Color scale shows the log number of frames in each bin, normalized to the total number of frames. **F.** Schematic illustrating firing rate predictions of the recruitment hierarchy. For example, if the fast neuron spikes, both intermediate and slow neurons should be firing (blue dotted line). **G.** Left) Spike rate of intermediate neuron in 30 ms preceding fast EMG spike (n = 4 pairs); center) slow neuron spike rate preceding fast EMG spike (n = 2 pairs); right) and slow neuron spike rate preceding intermediate EMG spike (n = 3 pairs).

We first compared force and velocity production in the 25 ms following each motor neuron spike (**Figure 5D, E**). Spikes in each motor neuron type tended to occur during specific regimes of force and velocity. Plots of probe position and movement during example epochs (**Figure 5D**) illustrate that fast motor neurons typically spiked when the tibia was already flexed and moving. In comparison, the only period during which slow motor neurons were silent was when the tibia was extending (negative probe velocity) or fully extended (zero probe position). As a result, the spike-triggered distribution of probe dynamics for the slow motor neuron is broad (**Figure 5E**, right). Large spike rate modulations in the slow motor neuron often coincided with gradual changes in the position of the probe, consistent with the slow kinetics of force production measured using current injection (**Figure 4**). In comparison, spikes in fast motor neurons caused the probe to rapidly accelerate and approach maximum velocity (**Figure 5D, E**). Intermediate neuron spikes could also produce small but measurable increases in force (**Figure 5B**, inset).

Overall, this analysis shows that each motor neuron type has a different regime of force probe position and velocity in which it is likely to spike (**Figure S5**). These regimes are overlapping. For example, in the regime where fast motor neuron spikes are likely, both intermediate and slow spikes are likely as well. Similarly, the slow motor neuron is likely to be spiking throughout the intermediate neuron’s regime. These relationships between motor neuron spiking and force production are consistent with the existence of a hierarchical recruitment order (**Figure 5F)**.

We tested this model by examining paired recordings of somatic (whole-cell) and axonal (EMG) spikes from different motor neurons. Neurons lower in the recruitment hierarchy were consistently active when a neuron producing more force was recruited (**Figure 5G).** In other words, activity in slow neurons preceded that of intermediate neurons, and activity in intermediate neurons preceded that of fast neurons. We rarely observed a violation of this recruitment order (**Figure S5**). The exceptions occurred when the fly was rapidly waving its leg, rather than pulling on the force probe. This situation is similar to rapid paw shaking behavior in cats, where recruitment order violations have also been observed (Smith et al., 1980). Interestingly, the membrane potential of each tibia flexor motor neuron reflected probe movement, even during periods of subthreshold activity (**Figure 5A-C**). This suggests that distinct motor neurons receive common synaptic input, and that recruitment order is determined by differences in spike threshold.

### A gradient of sensory input to motor neurons

So far, we have shown that tibia flexor motor neurons exhibit a gradient of functional properties (**Figures 2-4**) and a recruitment order during natural leg movements (**Figure 5**) that are both consistent with a size principle. As a further test of the size principle and recruitment order, we measured motor neuron responses to proprioceptive sensory feedback. A key function of proprioceptive feedback is to maintain stability by counteracting sudden perturbations, such as when an animal stumbles during walking (Tuthill and Azim, 2018). Larger perturbations require more corrective force, so as feedback increases it should result in the recruitment of additional motor neurons with increasing force production capacity.

We first compared the amplitude and dynamics of proprioceptive feedback in flexor motor neurons in response to ramping movements of the tibia (**Figure 6A-B**). Passive extension of the tibia caused excitatory postsynaptic potentials (PSPs) in all three flexor motor neurons, a response known as a resistance reflex. PSP amplitude was largest in slow motor neurons and smallest in fast neurons, as we might expect from observed differences in input resistance (**Figure 2**). In the fast and intermediate neurons, sensory-evoked PSPs typically failed to elicit spikes, whereas the firing rate of slow motor neurons was significantly modulated. Fast extension movements were sufficient to evoke single spikes in intermediate neurons (**Figure S6D-E**), but we never observed feedback-evoked spikes in fast motor neurons.

**Figure 6.**
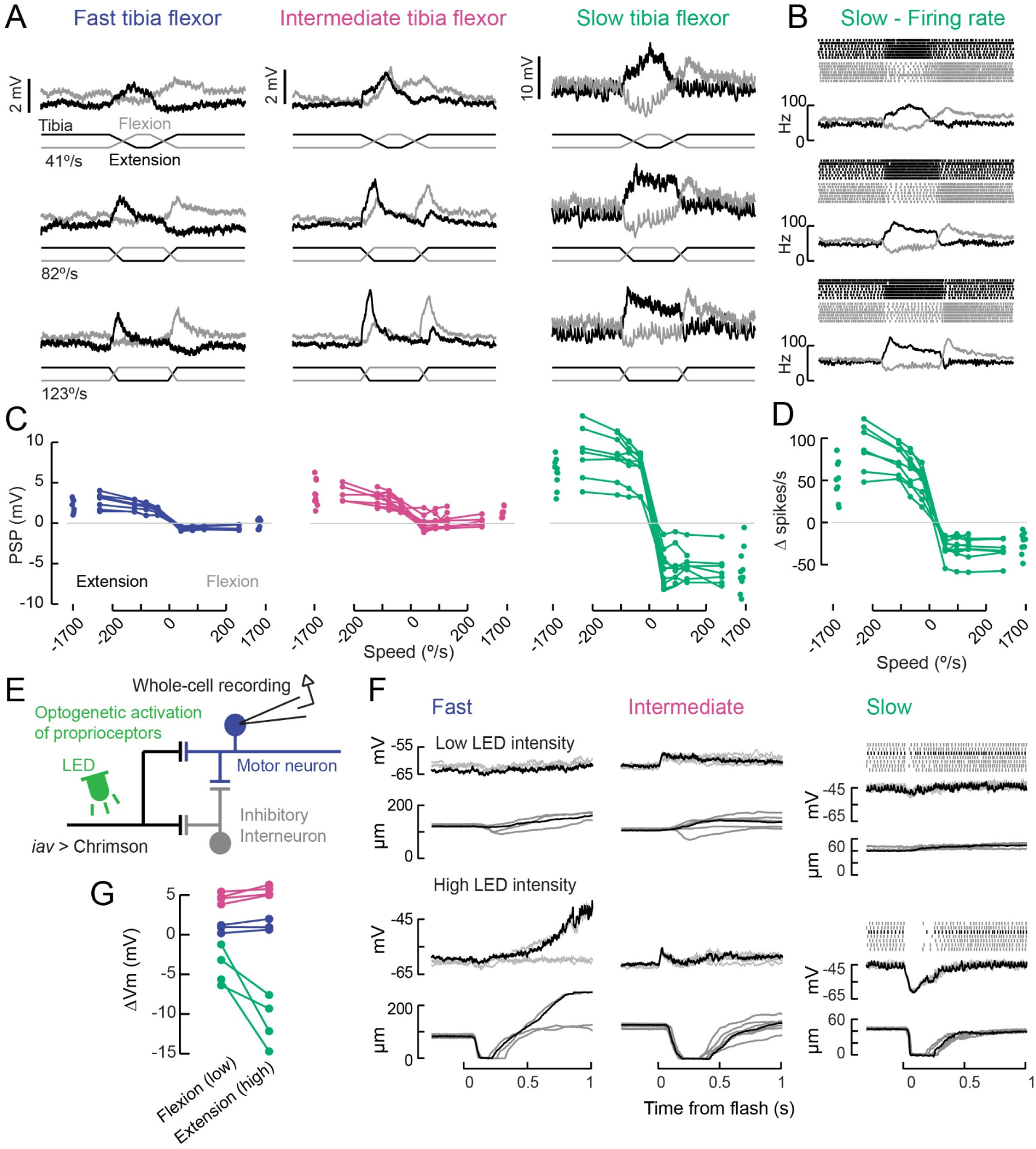
A gradient of proprioceptive feedback to motor neurons controlling tibia flexion. **A.** Whole-cell recordings of fast, intermediate, and slow motor neurons in response to flexion (black) and extension (gray) of the tibia. Each trace shows the average membrane potential of a single neuron (n = 7 trials). A 60 μm movement of the probe resulted in an 8° change in femur-tibia angle. **B.** Spike rasters and firing rates for the same slow neuron as in A. **C.** Amplitude of postsynaptic potentials vs. angular velocity of the femur-tibia angle (n = 7 fast, 7 intermediate, 9 slow cells). Responses to fast extension (−123°/s) are not significantly different for fast and intermediate neurons (p = 0.5) but are different for slow neurons (p<10^-5^, 2-way ANOVA, Tukey-Kramer correction for multiple comparisons). **D.** Amplitude of firing rate changes in slow motor neurons vs. angular velocity of the femur-tibia angle. **E.** Optogenetic activation of proprioceptive feedback to tibia motor neurons. The axons of femoral chordotonal neurons, labeled by *iav-LexA*, were stimulated with Chrimson activation during whole-cell recordings. **F.** Example recordings showing membrane potential, spiking activity, and probe position following an LED flash. Top: low intensity flashes caused the fly to mainly pull on the force probe, from individual fast (5.5 mW/mm^2^), intermediate (4.8 mW/mm^2^), and slow neurons (1.3 mW/mm^2^). Each trace is a single trial; the black traces belong to the same trial. Bottom: higher LED intensities caused the fly to let go of the force probe in fast (10.9 mW/mm^2^), intermediate (7.8 mW/mm^2^), and slow neurons (6.3 mW/mm^2^). **G.** Summary of optogenetically-evoked membrane potential changes in motor neurons for low (flexion-producing) and high (extension-producing) LED intensities. Fast and intermediate neurons depolarized initially, so the amplitude of the depolarization is quantified. Slow motor neurons exhibited stronger hyperpolarization.

Responses to tibia flexion were more diverse across the different motor neurons. Fast and slow motor neurons both hyperpolarized at flexion onset, while intermediate neurons depolarized with a longer latency. Sensory responses in fast, intermediate, and slow neurons were all similarly tuned across stimulus speeds (**Figure 6D-E**), amplitudes, and joint angles (**Figure S6A**). As reported in recordings from other insect species (Bässler and Büschges, 1998), we occasionally observed a reflex reversal, in which the sign of the proprioceptive PSP reversed (**Figure S6C**). In most instances, tibia extension led to excitatory responses in flexor motor neurons. But occasionally, typically when the fly was actively moving as the stimulus was delivered, extension produced an inhibitory PSP. Reflex reversal is an important mechanism that could allow the fly to switch between active and passive motor states (Bässler and Büschges, 1998; Tuthill and Wilson, 2016).

A major source of proprioceptive information about tibia position is the femoral chordotonal organ (FeCO). The *Drosophila* FeCO is comprised of ∼150 mechanosensory neurons that encode position, movement, and vibration of the femur-tibia joint (Mamiya et al., 2018). We used optogenetic activation (*iav-LexA*>*LexAOp-Chrimson*) to measure proprioceptive feedback from the FeCO to each motor neuron type (**Figure 6E**). In interpreting these experiments, it is important to note that Chrimson expression in FeCO neurons reduced motor neuron responses to passive leg movements compared to control flies (**Figure S6F**). We speculate that this side-effect of Chrimson expression is caused by a homeostatic decrease in the gain of proprioceptive feedback pathways following increased excitability.

Weak (low LED intensity) stimulation of FeCO axons caused the fly to gently flex its tibia, while higher intensity stimulation caused the fly to extend and then flex its tibia in a rapid and stereotyped manner (**Figure 6F**). As we observed with passive sensory stimulation (**Figure 6A-D**), FeCO activation had the largest effect on the membrane potential and spiking activity of slow motor neurons, and the weakest effect on fast motor neurons (**Figure 6G**). In slow neurons, these responses were hyperpolarizing, and grew stronger with increasing intensity of FeCO stimulation (**Figure 6F-G**). Responses in fast and intermediate neurons were typically depolarizing and did not vary as widely as a function of stimulation intensity.

Overall, these data demonstrate that the FeCO provides proprioceptive feedback to tibia flexor motor neurons, and that the amplitude and dynamics of proprioceptive feedback signals vary across the different motor neurons (**Figure 6A-D**). This gradient of sensory feedback amplitude is likely shaped by the concurrent gradient of motor neuron intrinsic properties (**Figure 3**). Although feedback signals were largely similar across motor neurons, we observed some differences in the sign and timing of proprioceptive signals that cannot be explained by the size principle. These differences suggest that proprioceptive feedback signals may be specialized for controlling particular motor neurons.

### Optogenetic perturbation of single motor neurons alters walking behavior

Our measurements of force production and recruitment order suggest that slow, intermediate, and fast tibia flexor motor neurons operate during distinct motor regimes. To test the implications of this organization for behavior, we used optogenetics to alter motor neuron activity in tethered, walking flies. We focused a green laser at the ventral thorax, at the base of the left front (T1) leg to optogenetically activate (*Gal4*>*UAS-Chrimson*, Klapoetke et al., 2014) or silence (*Gal4*>*UAS-gtACR1*, Mohammad et al., 2017) each motor neuron type. As a control, we used an “empty” Gal4 driver line, which has the same genetic background but lacks Gal4 expression.

To verify our optogenetic manipulations and characterize basic leg reflexes, we first measured the movement of the femur-tibia joint in headless flies with their legs unloaded (i.e., the fly was suspended in air, **Figure 7A**). Activation of fast motor neurons caused rapid flexion of the tibia (**Figure 7B, C, Movie S3**). We observed repeated patterns of tibia flexion and extension, suggesting that the fast neuron fired single spikes and that the resulting flexion was immediately countered by a resistance reflex. Activation of intermediate neurons caused slower tibia flexion, while activation of slow neurons caused even slower contractions, often followed by prolonged tibia extension. Silencing each motor neuron type did not consistently evoke leg movements, though silencing the full motor neuron population resulted in paralysis (**Movie S4**). These results are consistent with our measures of force production during electrophysiology experiments (**Figures 4, 5**) and provide a useful baseline for interpreting results from walking flies.

**Figure 7.**
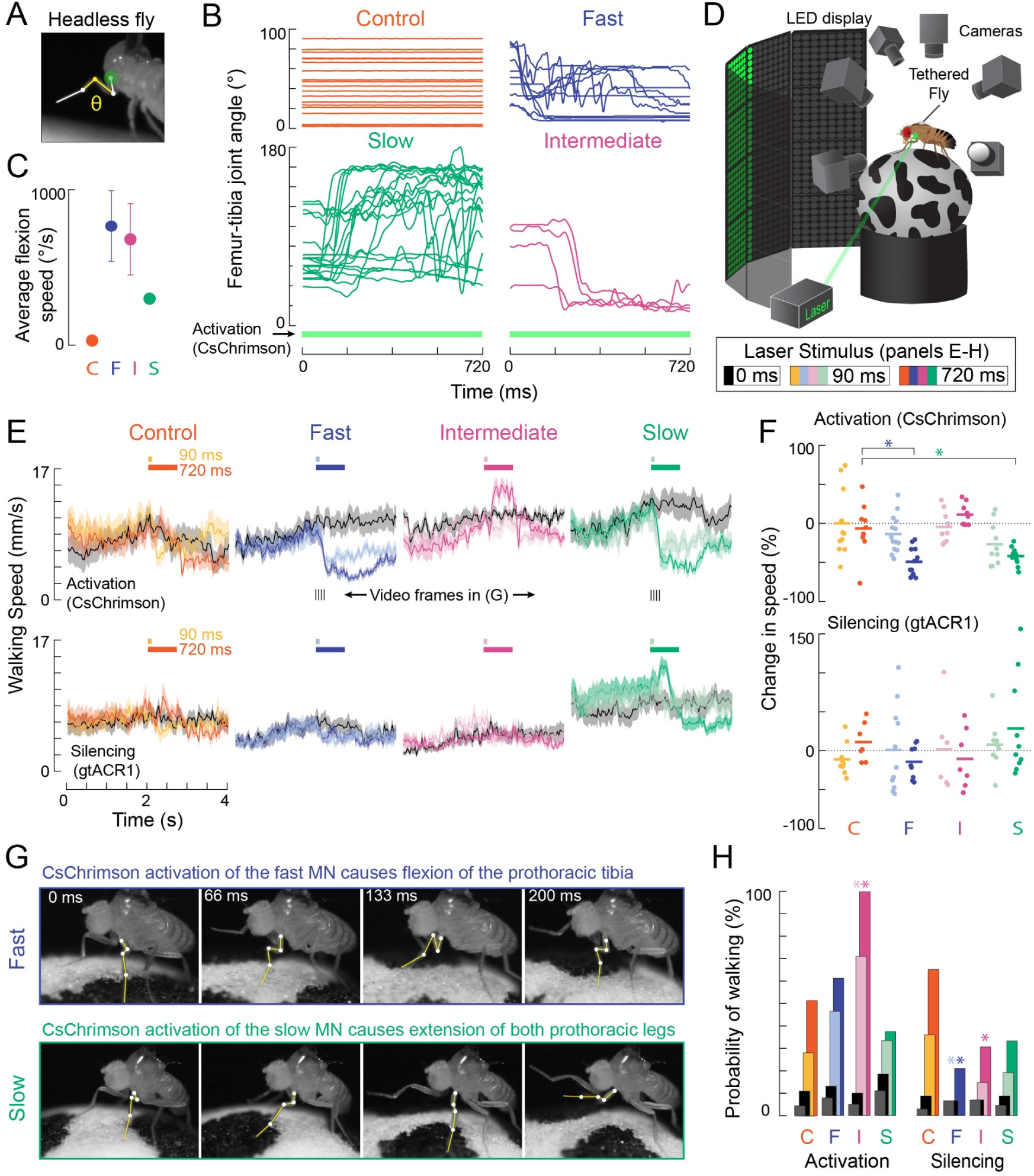
Optogenetic perturbation of motor neurons in behaving flies. **A.** Example frame illustrating behavioral effects of optogenetically activating leg motor neurons in headless flies. A green laser (530 nm) is focused at the coxa-body joint of the fly’s left front leg (outlined in white) and the femur-tibia joint (yellow) is monitored with high-speed video. **B.** Example traces of femur-tibia joint angles during optogenetic activation (*Gal4>csChrimson*) of each motor neuron type in headless flies. The control line has the same genetic background as motor neuron lines but lacks Gal4 expression (*BDP-Gal4>csChrimson*). **C.** Average (±sem) tibia flexion speed during the first 300 ms of csChrimson activation (n=5 flies of each genotype). **D.** Schematic of the behavioral setup, in which a tethered fly walks on a spherical treadmill. The treadmill and fly are tracked with separate cameras. As in **A**, a green laser is focused on the coxa of the left front leg. **E.** Average treadmill forward velocity (±sem) of walking flies, when activating (top row) or silencing (bottom row) control (n=11, 13), fast (n=15, 14), intermediate (n=9,12), or slow (n=11, 13) motor neurons for 90 ms (dark colors) or 720 ms (light colors). Black traces indicate trials with no laser (0 ms). Grey boxes indicate the period of optogenetic stimulation. Black dashes above fast and slow flexor activation traces indicate the time points in panel G. **F.** Percent change (change in speed/pre-stimulus speed) in the average running speed of flies shown in C during stimulation+200 ms for activation (top) and silencing (bottom). Asterisks indicate p<0.01 relative to control (bootstrapping with false discovery rate correction; see Experimental Procedures for details). **G.** Example frames showing representative behavioral responses of running flies during the first 200 ms of fast (top) and slow (bottom) MN activation. **H.** Probability that a stationary fly initiated a walking bout during the 500 ms following either MN activation (left) or silencing (right). Dark colors are for the 90 ms stimulus (or no stimulus – black) and light colors are for the 720 ms stimulus (or no stimulus – grey). Asterisks indicate p<0.01 relative to control (bootstrapping with false discovery rate correction; see Experimental Procedures for details).

To optogenetically manipulate motor neuron activity during walking, we positioned tethered flies on a spherical treadmill within a visual arena (Reiser and Dickinson, 2008; **Figure 7D**). Previous studies have found that walking speed and gait on the treadmill are similar to freely walking flies (Szczecinski et al., 2018). In our setup, the fly was motivated to walk forward with a wiggling stripe (**Figure 7E**), and we tracked the treadmill and the fly’s behavior with multiple high-speed cameras.

We first asked how flies would respond to transient optogenetic activation of the different tibia flexor motor neurons. Activation of the fast motor neuron caused walking flies to slow down, as measured by a decrease in forward ball velocity. In videos of leg movements, flies appeared to slow because the tibia of the front leg flexed and did not extend again, thus interrupting walking (**Figure 7E, Movie S5)**. Activation of the slow motor neurons also interrupted walking (**Figure 7E**), but this was because the fly reacted by extending its tibia, as when the legs were unloaded (**Movie S7)**. Based on the muscle fibers innervated by the slow motor neuron, we speculate that optogenetic activation of this neuron may increase twisting at the joint, which the fly reacts to with a compensatory extension of the leg. Unlike the other two cells, activation of the intermediate motor neuron appeared to cause the flies to transiently increase their walking speed. Activation of the intermediate motor neuron in stationary flies also caused them to start walking (**Figure 7E-F, Movie S6**). Extra spikes in the intermediate tibia flexor motor neuron may accelerate the rotation of the ball, forcing the other legs to move faster in response. Overall, these results show that activation of different motor neurons drive distinct behavioral responses that depend on the fly’s behavioral state (e.g., walking vs. standing).

Optogenetically silencing motor neurons had less pronounced effects on walking speed (**Figure 7E**). However, silencing of fast and intermediate neurons decreased the probability that a stationary fly would initiate walking compared to controls (**Figure 7H, Movies S6 and S7**). These results suggest that motor neurons with a high force production capacity (**Figure 4**) may be dispensable for normal walking behavior, but their activity may be important for producing the high levels of force required to initiate walking.

## Discussion

In this study, we describe a gradient of cellular morphology, electrical excitability, force production, and proprioceptive feedback among motor neurons that control flexion of the *Drosophila* tibia. At one end of the spectrum is the slow motor neuron, which has fine axons and dendrites, high input resistance, a high spontaneous firing rate, strong sensory input, and produces low force per spike. At the other end, the fast motor neuron has larger diameter axons and dendrites, low input resistance, a low spontaneous firing rate, weak sensory input, and produces large forces with just a single spike. These properties are consistent with relationships first described among vertebrate motor neurons, collectively known as the size principle (Henneman and Olson, 1965; Henneman et al., 1965b; Kernell, 2006; Mcphedran et al., 1965; Wuerker et al., 1965).

A second, and related, organizing principle of vertebrate motor control is the recruitment order. Motor neurons within a pool tend to fire in a temporal sequence: small, low gain motor neurons that generate weak forces are recruited first, and larger, more powerful motor neurons are recruited later (Henneman, 1957; Henneman et al., 1965a; Mendell, 2005). We observed a similar relationship among tibia flexor muscles and motor neurons in the fly leg. Activity in the fast motor neuron was preceded by activity in slow and intermediate motor neurons. In paired recordings between flexor motor neurons, we rarely observed violations of this recruitment hierarchy.

Together, the size principle and recruitment hierarchy suggest a simple mechanism for graded control of joint movement: increasing the synaptic input to the motor neuron pool 1) increases the spike rates of neurons currently recruited, and 2) recruits higher gain motor units. As a result, recruitment of more motor units increases force nonlinearly, overcoming suppressive nonlinearities in spike rates and muscle force production (Bakels and Kernell, 1994; Monster and Chan, 1977). Furthermore, the relative increase in force does not decrease with successive recruitment, as it would if all motor units produced similar amounts of force. Thus, much like Weber’s law describes the constant sensitivity to relative stimulus intensity (i.e. contrast; Fechner, 1860), a recruitment hierarchy based on the size principle maximizes the resolution of motor unit force while also simplifying the dimensionality of the motor system (Henneman et al., 1965a; Mendell, 2005).

### Organization of leg motor pools

The muscles that control flexion of the *Drosophila* tibia are innervated by at least 15 motor neurons (Baek and Mann, 2009; Brierley et al., 2012). Here, we chose to analyze three identified neurons that represent the extremes of this motor pool. The fast tibia flexor labeled by *R81A07-Gal4* innervates the entire distal compartment of the flexor muscle and produces the largest extracellular spikes we recorded in the leg. The slow femur flexor labeled by *R35C09-Gal4* innervates a small muscle located at the distal tip of the femur. Recordings from motor neurons innervating slightly more proximal flexor muscle fibers produced more force per spike (**Figure S4**) indicating that the *R35C09-Gal4* neuron is among the weakest. Finally, we studied an intermediate motor neuron labeled by *R22A08-Gal4* which innervates separate muscle fibers from the fast and slow neurons and occupies an intermediate functional regime. Each of these motor neurons was stereotyped across flies in both its anatomy and function. Untargeted recordings from other femur flexor motor neurons (**Figure S4**) were also consistent with the gradient of properties we describe. Those data suggest that morphological and functional relationships hold for other members of the motor pool, such that the slow and intermediate neurons are representative of more coarsely-grouped “slow” and “intermediate” cell types.

Some of the properties we measured were intrinsic to the motor neurons themselves, like their morphology and input resistance, while others could arise from the properties of other circuit elements (e.g., muscle fibers and presynaptic neurons). For example, the gradient in resting membrane potential likely reflects differences in sustained synaptic input that establish the resting flexion force (**Figures 4 and S4**). The gradients in functional properties correlate with the developmental birth order of leg motor neurons: the fast tibia flexor is born embryonically, and then more proximal-targeting neurons are born before neurons with distal axonal projections (Baek and Mann, 2009; Brierley et al., 2012). Thus, the principles governing motor neuron development (Venkatasubramanian et al., 2019) and circuit assembly (Enriquez et al., 2015) may also determine their physiological properties.

The structure of the leg motor system in *Drosophila* is in some ways similar to other well-studied walking insects. In the metathoracic leg of the locust, a single tibia flexor muscle is innervated by nine motor neurons that were also classified into three groups: fast, intermediate, and slow (Burrows and Hoyle, 1973; Phillips, 1981; Sasaki and Burrows, 1998). Different motor neurons within this pool lie along a gradient of intrinsic properties (Sasaki and Burrows, 1998) and are sensitive to different types of proprioceptive feedback, such as position vs. velocity or fast vs. slow movements (Field and Burrows, 1982; Newland and Kondoh, 1997). The femur of the stick insect is innervated by flexor motor neurons that were also described as slow, semi-fast, and fast, based on their intrinsic properties and firing patterns during behavior (Bässler, 1993; Schmidt et al., 2001).

Much of the work in bigger insects and crustaceans has focused on the most reliably identifiable neurons, the fast and slow extensor tibiae (FETi and SETi), antagonists to the more diverse tibia flexor motor neurons. Consequently, details of how sensory feedback and local interneurons recruit specific subsets of flexor motor neurons has been relatively understudied (Clarac et al., 2000) (but see (Gabriel and Büschges, 2007; Gabriel et al., 2003; Hill and Cattaert, 2008)). Our work exemplifies the advantage of using *Drosophila* genetics to identify cell types, and even individual cells, within a diverse motor pool. Genetic markers have also recently been identified for tibia extensor motor neurons in flies (Venkatasubramanian et al., 2019), which will enable future investigation of premotor input to antagonist motor neurons.

Flies differ from these other invertebrates in one major respect: while most arthropods possess GABAergic motor neurons that directly inhibit leg muscles, holometabolous insects such as *Drosophila* do not (Schmid et al., 1999; Witten and Truman, 1998). The presence of inhibitory motor neurons has been proposed as a key underlying reason why insects have been able to achieve flexible motor control with small numbers of motor neurons (Belanger, 2005; Wolf, 2014). That flies lack this capability means that other mechanisms must be at play. Fly legs are innervated by neurons that release neuromodulators, such as octopamine, which could underlie flexible tuning of muscle excitability (Zumstein, 2004).

### Function of distinct leg motor neurons

The large range of gain, or force production, at the femur-tibia joint implies that different subgroups of flexor neurons will be recruited during distinct behaviors. Activity in slow motor neurons produces small, low force movements, while fast motor neuron activity produces fast, high amplitude movements. Consistent with this division, the output of slow motor neurons is more strongly influenced by feedback from leg proprioceptors (**Figure 6**). Thus, proprioceptive feedback may continuously modulate the firing rate of slow motor neurons for precise stabilization of posture during standing or grooming. When the slow motor neuron is optogenetically activated (**Figure 7**), flies stop walking and extend their legs. The reason for this response is unclear but could reflect the fly’s reaction to a loss of autonomous motor control.

In contrast to the postural movements controlled by slow motor neurons, rapid, stereotyped movements like escape (Card and Dickinson, 2008; Zumstein, 2004) are likely to use fast motor neurons whose activity is less dependent on sensory feedback. This division of labor may also provide energy efficiency: slow contracting muscle fibers use aerobic metabolism and take advantage of energy stores in the form of glycogen, while fast muscle fibers are anaerobic and lack energy stores, leading to more rapid fatigue (Kernell, 2006).

Even for a specific behavior, such as walking, different motor neurons may be recruited in different environments and contexts. We saw that optogenetically activating intermediate motor neurons caused stationary flies to start walking and walking flies to walk faster (**Figure 7**). In a similar manner, stick insects walking on treadmills typically recruit fast motor neurons only during fast walking and, in that case, late in the stance phase. But as friction on the treadmill increases, fast neurons are recruited earlier during stance (Gabriel et al., 2003). Leg kinematics and force production also change as insects walk up or down inclines (Dallmann et al., 2019; Wöhrl et al., 2017). When locusts (Duch and Pflüger, 1995) and cockroaches (Larsen et al., 1995; Watson and Ritzmann, 1997; Watson and Ritzmann, 1998) are forced to walk upside-down, fast flexor neurons are recruited to allow the animal to grip the substrate. This context-dependence is not limited to walking: small changes in body posture can have a large effect on which motor neurons are recruited during target reaching movements in locusts (Page et al., 2008).

In this study, we chose to focus on flexion of the femur-tibia joint of the fly’s front leg. How might these results compare to motor neuron pools of antagonist muscles? Tibia flexor motor neurons outnumber the extensor neurons (Baek and Mann, 2009; Brierley et al., 2012), and thus are likely to possess a shallower gradient of intrinsic and functional properties, as has been found in crayfish (Hill and Cattaert, 2008). Invertebrate muscles also exhibit polyneural innervation, such that a given muscle fiber may be innervated by multiple motor neurons (Brierley et al., 2012; Hoyle, 1955; Sasaki and Burrows, 1998). Polyneural innervation could allow independent activation of motor neurons in order to use the same muscle fibers during different contexts, or it could make the force produced by one motor neuron dependent on coincident activity in another motor neuron (Sasaki and Burrows, 1998).

### The Importance of Spike Timing in Leg Motor Control

We found that single spikes in fast flexor motor neurons produce enough force to cause large changes in the femur-tibia joint angle. When the fly spontaneously pulled on the force probe, fast motor neuron activity was sparse, often consisting of isolated spikes (**Figures 1 and 5**). These results indicate that spike timing is a critical variable for motor control of the fly leg. The importance of motor neuron spike timing has recently been shown in other systems, including the vocal muscles of the songbird and the flight motor system of the hawkmoth (Sober et al., 2018).

Previous work has also shown that precisely when a muscle is activated can determine its function during different motor contexts (Dickinson et al., 2000). For example, depending on its activation relative to the changing muscle length, a muscle can actuate a limb or it can act as a brake upon movement generated by other muscles (Ahn and Full, 2002). When the fly is standing still, reflexes recruit antagonist muscles to counteract a sudden lengthening of the muscle (brake). Theses reflexes must be overridden when the fly begins to move and the same motor neuron now acts to shorten the muscle (Clarac et al., 2000). Indeed, we observed instances of reflex reversal when the fly actively moved its tibia (**Figure S6**), demonstrating that premotor circuits can change the way sensory feedback shapes the temporal patterns of motor activity.

### Organization of Premotor Circuits

The size principle proposes that a gradient of excitability coupled with a gradient in contractile force and speed provides a simple mechanism by which motor circuits adjust muscle force production and contraction timing. Increasing excitatory synaptic drive common to all members of the motor neuron pool could sequentially recruit neurons, and neurons recruited later in the hierarchy would be also more susceptible to a common source of inhibition and thus de-recruitment (Henneman et al., 1965a). The alternative is that premotor neurons have diverse cellular properties and synapse physiology, and are specifically connected to subsets of motor neurons in a pool. Much previous work has adopted the simplifying assumption that premotor input to a motor pool is comprised of cell types with homogeneous properties and connectivity (Grillner and Jessell, 2009).

While conceptually useful, this assumption is complicated by instances in which recruitment order can change in different contexts. For instance, muscles controlling human fingers can change recruitment order based on movement direction (Desmedt and Godaux, 1981). Another exception is rapid paw shaking behavior in cats, which relies exclusively on activity in the fast twitch gastrocnemius muscle, rather than on hierarchical recruitment together with the more heterogeneous soleus muscle (Smith et al., 1980). During rapid escape behaviors in zebrafish, slow motor neurons can completely drop out of the population firing pattern (Menelaou and McLean, 2012). These specific behaviors thus suggest that movement has more degrees of freedom than can be supported by recruitment of motor neurons based on their excitability alone. Although our results are consistent with a recruitment hierarchy among tibia flexor motor neurons, we anticipate that studies of this motor neuron pool during different contexts and behaviors would likely reveal exceptions. Indeed, in a small fraction of instances (∼50/3082), we observed that intermediate motor neuron spikes preceded slow motor neurons spikes while the fly rapidly moved its leg (**Figure S5**), likely due to inhibition of slow motor neuron firing.

Little is currently known about the premotor circuitry for limb coordination in *Drosophila*. In contrast, recent work in the vertebrate spinal cord has begun to reveal the modular organization of premotor circuits that control locomotion (Kiehn, 2016; McLean and Dougherty, 2015). For example, *in vivo* recordings in turtles and zebrafish have shown that motor neurons receive coincident excitation and inhibition (Berg et al., 2007; Kishore et al., 2014), which may selectively de-recruit motor neurons (Henneman et al., 1965a). Populations of premotor neurons also display gradients of electrical excitability and recruitment (Ampatzis et al., 2014; Song et al., 2018). Finally, premotor neurons form microcircuits (Callahan et al., 2019; Menelaou et al., 2019) that integrate feedback from motor neurons (Song et al., 2016) and can flexibly switch between different motor neuron recruitment patterns (Bagnall and McLean, 2014). Our characterization of motor neuron diversity provides a handle to investigate similar premotor circuit motifs for flexible limb motor control in the fly.

### Summary

In this study, we describe a simple organizing principle for how motor neurons control tibia movement in *Drosophila*. This principle provides a useful framework for understanding the organization and function of motor circuits that flexibly control a wide range of fly behaviors, from walking to aggression to courtship. Studies of motor control in the fly VNC are now possible thanks to an increasing catalog of genetically-identified cell types (Lacin et al., 2019; Shepherd et al., 2019, Namiki et al., 2018), connectomic reconstruction with serial-section EM (Maniates-Selvin et al., 2019) and the ability to image from VNC neurons in walking flies (Chen et al., 2018). We anticipate that *Drosophila* will provide a useful complement to other model organisms in understanding the neural basis of flexible motor control.

## Acknowledgements

We thank Jim Truman, Wei-Chung Allen Lee, Jasper Maniates-Selvin, Eiman Azim, Randy Powers, Marc Binder, and Brendan Lehnert, as well as members of the FlyLoops U19 team (Tom Clandinin, Michael Dickinson, Shaul Druckmann, Richard Murray, and Rachel Wilson) for helpful discussions, and members of the Tuthill laboratory for feedback on the manuscript. We thank Peter Detwiler, Fred Rieke, and Rachel Wong for generous sharing of equipment, Shellee Cunnington for preparation of solutions, Eric Martinson and Bryan Venema for technical assistance, and Michael Reiser and Michael Dickinson for sharing fly stocks. Stocks obtained from the Bloomington Drosophila Stock Center (NIH P40OD018537) were used in this study. We also acknowledge support from the NIH (S10 OD016240) to the Keck Imaging Center at UW, and the assistance of its manager, Nathaniel Peters. A.W.A. was supported by NIH fellowship F32 DC013928. This work was funded by a Searle Scholar Award, the McKnight Foundation, and NIH grant U19NS104655 to J.C.T.

## Author Contributions

A.W.A and J.C.T. conceived the project and designed the experiments. A.W.A performed calcium imaging and electrophysiology experiments and analyzed the data. P.G. performed immunohistochemistry experiments. E.D. conceived and performed behavioral experiments and analyzed the data. R.S.M. and L.V. assisted with identification of motor neuron Gal4 lines. A.W.A and J.C.T. wrote the paper.

## Experimental Procedures

### Identifying Gal4 lines that label leg motor neurons

Images of Gal4 expression in the VNC in the Janelia FlyLight collection (Jenett et al., 2012) were screened to find lines that sparsely labeled motor neurons. Flies were obtained from the BDSC, and we imaged leg expression of GFP to characterize muscle innervation patterns.

Our muscle nomenclature is based on (Soler, 2004), which also serves as the basis for the leg motor neuron nomenclature (Baek and Mann, 2009; Brierley et al., 2012). For clarity, however, we refer to muscles of the femur as flexors or extensors, rather than as depressors or levators (relative to the natural stance of the insect). Soler et al. in turn based their nomenclature on (Miller, 1950). There appears to be a discrepancy between the two: the muscle named the tibia reductor muscle by Soler et al. is described as one of two depressor muscles by Miller, his muscles 40 and 41. Miller applied the nomenclature of (R. E. Snodgrass, 1935) (tibia depressor muscles 136a and 136b) to *Drosophila* leg muscles. Muscles that control the tibia are located in the femur, etc, and Snodgrass described a reductor of the femur, located in the trochanter, perhaps leading to this misreading. The trochanter-femur joint in most insects has limited mobility. Presumably, Snodgrass named it a reductor muscle because depressor or levator would be inaccurate. We agree with the characterization of Miller and Soler that muscles 40 and 41 are two distinct muscles as they attach via distinct tendons, as seen in X-ray images of the leg musculature (Pacureanu et al., 2019). Because the function of muscle 41 is to flex the tibia, for clarity we refer to it here as a tibia flexor.

### Fly husbandry

*Drosophila* were raised on cornmeal agar food on a 14h dark/10h light cycle at 25°C and 70% relative humidity. Females flies, 1-4 days post eclosion, were used for all experiments except tethered behavior experiments. For tethered behavior experiments, both male and female dark-reared flies, between 2-10 days post-eclosion, were used. For experiments involving optogenetic reagents (Chrimson variants and gtACR1), adult flies were placed on cornmeal agar with all-trans-retinal (35mM in 95% EtOH, Santa Cruz Biotechnology) for 24 hrs prior to the experiment. Vials were wrapped in foil to reduce optogenetic activation during development.

### Electrophysiology and calcium imaging preparation

Flies were positioned in a custom steel holder as described in (Tuthill and Wilson, 2016), with modifications to allow us to image the movements of the fly leg. Flies were anesthetized on ice <2 min, so that they could be positioned ventral side up, with head and thorax fixed in place with UV-cured glue. The front legs were glued to the horizontal top of the holder, the coxa aligned with the thorax, and the femur positioned at a right angle to the coxa and body axis. In this configuration, the fly could wave its tibia freely in an arc at an angle of ∼50-65° to the top surface of the holder. The holder was placed in the imaging plane of a Sutter SOM moveable objective microscope. All recordings were performed in extracellular fly saline (recipe below) at room temperature.

A water immersion 40X objective (Nikon) was used for whole-cell patch-clamp experiments and placing electrodes in the leg. We used a 5X objective (Nikon) to view the fly’s right front femur and tibia through the saline during spontaneous movements of the leg. Videos of the preparation were acquired at 170 Hz through the 5X objective with a Basler acA1300-200um machine vision camera. Custom acquisition code written in MATLAB (https://github.com/tony-azevedo/FlySound) controlled generation and acquisition of digital and analog signals through a DAQ (National Instruments). Input signals were digitized at 50 kHz.

### Force measurement and spring constant calibration

To measure forces produced by leg movements, we recorded video of the position of a flexible “force probe” as the fly pulled against it. The force probe was a PBT filament fiber from a synthetic paint brush (Proform CS2.5 AS), threaded through the end of a glass micropipette (1.5 mm OD, 1.1 mm ID, WPI). UV-cured glue was sucked up into the micropipette, the fiber was threaded through, leaving ∼1-2 cm out from the tip of the glass, and the glue was cured. The micropipette allowed us to mount the force probe in a custom holder and to couple it to a micromanipulator (Sutter MP-285). Videos of the force probe were acquired at 170 Hz through the 5X objective with a Basler acA1300-200um machine vision camera. One pixel equaled 1.03×1.03 μm^2^. We wrote custom machine vision code that detects the position of the force probe in each frame of the video by 1) allowing the user to draw a line along the probe in the video, 2) rotating the image of the probe perpendicular to the line, 3) averaging the lines of the video into a single line with a hump of intensity where the out-of-focus probe was, and then 4) finding the center of mass of the hump of intensity, ±FWHM above baseline. A similar technique employing a probe to measure force has been used in *Drosophila* in previous studies (Elliott and Sparrow, 2012).

At steady state, the position of the force probe was related to the force through a spring constant, k, F = −kx (Figure S2). We measured the spring constant by positioning the force probe over an analytical balance. A glass coverslip was oriented vertically on edge in a piece of wax stuck onto the balance, and the tip of the force probe was positioned at the top edge of the coverslip. We then moved the micromanipulator to different positions. The “mass” of the force probe was multiplied by gravity to give the force at that position. We then fit a linear relationship between force and position to measure the spring constant (Figure S2A). The force probe we used for experiments in this study had a spring constant of k = 0.2234 µN/µm, and stuck out approximately 1.5 cm past the end of the micropipette.

The force probe was not only a spring, but also had mass and was placed in saline, so inertia and drag affected its dynamics, which we modeled as *F* = *m d*^2^*x*/*dt*^2^ + *c dx*/*dt* + *kx*. To measure these properties, we “flicked” the force probe by moving it to different positions with a glass micropipette, abruptly letting go, and allowing the probe to relax back to rest (Figure S2B). We imaged the position of the probe at 1.2 kHz, using a restricted region of interest on the camera, and extracted dynamical parameters m = 0.1702 mg and c = 0.1377 kg/s. While not zero, the effective mass and drag were negligible (Figure S2C). The probe was slightly under-damped with a relaxation time constant of 2.5 ms and oscillation period of 5.8 ms, such that when imaging spontaneous movements at 170 Hz, the probe would effectively come to rest within 1 frame. The relaxation time constant was much faster than the fly’s spontaneous movements, even when the fly was attempting to let go of the force probe.

In Figure 4D-F, we calculate force by including drag and inertia, but in other figures we report leg displacement and the approximation of force, assuming that drag and inertia are negligible. We easily captured the lateral movement of the force probe across the frame but avoided estimating the vertical movement as the fly pulled the probe closer to its leg. As a result, the displacement of the force probe, and thus the force, that we measure in Figures 1 and 5 is underestimated.

### Mechanical stimulation of the leg

To move the leg and passively stimulate sensory feedback, we mounted the force probe perpendicularly on a piezoelectric actuator with a 60 μm travel range (Physik Instrumente) and a strain gauge. The axial position of the probe was controlled by an amplifier (Physik), with voltage commands generated in MATLAB and delivered through the DAQ board (National Instruments). The output of the actuator’s strain gauge was used to control the position of the actuator through closed-loop feedback. The strain gauge sensor output was sampled at 50 kHz. The probe tip was positioned near the end of the tibia, giving a lever arm of 417±7 (s.d.) μm across flies (n=8). We then moved the leg through its arc of rotation until it was approximately at 90°. To measure the effect of leg angle on the amplitude of sensory feedback, we then moved the probe to a range of axial positions (−150 μm = −21°, −75 μm = −10°, 75 μm = 10° and 150 μm = 21°, negative direction is extension) and repeated the stimuli (Figure S6A).

We delivered ramp stimuli that moved the leg 60 μm = 8° with varying speeds, in both flexion (+) and extension (−) directions. We measured the actual speed of step stimuli by finding the maximum derivative of the strain gauge signal during the step onset. The range of angular velocities produced by the probe span the range shown to activate position and velocity sensitive femoral chordotonal neurons in the femur (Mamiya et al., 2018). The force probe was flexible, so the actual displacement of the leg could be affected by passive forces of the joint as well as recruitment of motor neurons. Indeed, when we imaged the position of the force probe during the mechanical stimuli, the position departed from the desired position, e.g. by <3 μm or <0.8° for 8° ramp stimuli. The errors in displacement of the force probe were small, so we did not attempt to correct them by, for instance, using a rigid probe instead of the flexible probe.

To generate larger mechanical perturbations than we could deliver with the probe, we used a glass hook to draw the tip of the flexible force probe away from the tibia, loading the force probe. We then released the tip of the probe by continuing to draw the hook away until it lost contact with the probe, causing the probe to flick back and whack the leg (Figure S6).

### Leg tracking

In trials where the fly was free to wave its leg rather than pull on the force probe, we tracked the leg using DeepLabCut (Mathis et al., 2018). For a training set, we labeled the tibia position for ∼45 frames for 3 different videos from each fly using custom labeling code (https://github.com/tuthill-lab/LabelSelectedFramesForDLC). We labeled 6 points on the stationary femur, 6 points on the tibia, as well as prominent bright objects that would otherwise often be falsely identified as part of the legs, including the EMG electrode, the force probe, and several specular creases in the steel holder. The resnet50 network used in DeepLabCut served as the starting point for training, but as we added more flies to the training set, we initiated further training from the previously trained network. We found that the networks failed to generalize across flies but that ∼150 labeled frames were sufficient to ensure >99% accuracy on other frames for that fly (Figure S1B).

In post-processing, we measured the distribution of pairwise nearest neighbor distances between the 6 detected tibia points and assumed that outliers indicated that a point was poorly identified. If only a single point was misidentified (∼0.7 %), we filled in the point with random draws from the nearest-neighbor pairwise distance distributions. The network misidentified more than one point 0.2% of the time, typically when the fly moved its leg particularly quickly, blurring the image across several frames, which we then excluded. We median-filtered the x, y coordinates for each tibia point across frames in the video, and found the centroid of the 6 points, approximately the middle of the tibia. These centroid points traced out an ellipse that was the projection of the circular arc of the leg in the plane of the camera. Fitting an ellipse to the centroids allowed us to calculate the azimuthal angle of the leg arc (∼50-65°) and the real angle between the stationary femur and the moving leg. We then used the real angle of the leg to detect when the leg was extended (>120°) or flexed (<30°) (Figure S1E).

### Calcium imaging

To link the activation of muscle fibers to the movement of the leg or the force probe, we imaged the calcium influx into muscles with GCaMP (Figure 1, Figure S1A). We used MHC-LexA to drive expression of GCaMP6f in muscles. Epifluorescent 488 nm illumination excited GCaMP fluorescence. As above, IR light, reflected off the leg and force probe, entered the objective together with GCaMP fluorescence. IR light passed through a second long-pass dichroic (560 nm, Semrock) to the leg imaging camera while GCaMP wavelengths were reflected into a second Basler camera imaging at 50-60 Hz. The imaging window of the GCaMP camera was restricted to the femur and the video frames were registered (Guizar-Sicairos et al., 2008) to remove vibrations due to movements of the saline meniscus (Movie S1).

In the dark, the fly tended not to move its leg. However, as soon as the blue epifluorescent light turned on to excite the calcium sensor, the fly began to struggle and move its leg. We imaged spontaneous movements under two conditions. 1) The fly could pull on the force probe with its tibia or 2) the fly waved its leg around spontaneously without the force probe. To make sure the tibia did not obscure the calcium signals in the femur, a glass hook was placed near the femur as a barrier to prevent the fly from completely flexing the leg.

### Clustering of calcium signals

To measure which portions of the leg musculature contracted together, we used the k-means algorithm in MATLAB to calculate clusters from calcium signals imaged at 50 Hz while the fly waved its leg with no force probe. We drew an ROI around the femur, excluding only points near the femur tibia joint where intensity changes were dominated by the tibia obscuring the femur. We calculated k=3-8 clusters, with the correlation of pixel intensity as the distance metric. Once the pixels were assigned to a cluster, we applied a Gaussian kernel to the pixel cluster assignments and excluded pixels which fell below 0.75, indicating that less than ¾ of the surrounding pixels were of the same cluster. This resulted in clearly defined clusters with gaps between them but no major gaps, indicating that similar clusters were also grouped anatomically (Figure S1C). To confirm that the clustering indicated changes in calcium influx rather than muscle movement, we also ran the same clustering routines on flies expressing GFP in the muscles (Figure S1G-H). In this case, clusters were dispersed, fluctuated very little in intensity and bore no similarity to musculature.

We found that with k = 6, the clusters were broadly similar across flies. We numbered the 6 clusters as follows: Cluster 1 was large and distal/ventral; Cluster 2 was proximal and ventral; Cluster 3 ran down the center of the leg, neighboring Cluster 2; Clusters 4, 5, 6 were assigned from proximal to distal. With fewer than 5 clusters, the proximal cluster tended to be much larger, incorporating much of the region labeled as cluster 3 (Figure S1F). With more than 6 clusters, the smallest and least modulated clusters tended to divide, not giving us any further information. k=6 clusters potentially allowed for pixels that were most correlated with extension of the leg to cluster together, but we did not see large increases in fluorescence with extension.

The pixels assigned to each cluster were averaged to give a cluster intensity for each frame. For trials in which the fly pulled on the force probe and the force probe could obscure the femur, we used the same clusters calculated above but used smaller ROI to include only the proximal portion of the leg. If the force probe still happened to obscure some of the pixels (>40%), that frame was excluded from the analysis.

The time constant of the calcium indicator was slower than the fly’s movements, such that fluorescence built up over subsequent contractions and pixel intensity (ΔF/F) was not directly related to contraction of the muscle. Instead, we took positive increases in cluster intensity to indicate muscle activation and neural activity. The calcium signals were imaged at 50 Hz to improve pixel signal to noise, whereas the leg and force probe position were imaged at 170 Hz. To align the samples, we applied a Sovitzky-Golay interpolation to the cluster calcium signals, using the derivative of the local spline as the derivative of the ΔF/F for each cluster (sgolay_t function in MATLAB, by Tiago Ramos, N=7, F=9). Negative derivatives reflected noise in the cluster intensities, so positive rectified derivatives were then thresholded above 2 standard deviations of the noise to indicate cluster “activations”.

Surprisingly, we did not identify any clusters that increased their calcium activity during tibia extension. Flies occasionally held their legs extended (Movie S1), at which point we expected to see some clusters increase fluorescence (Figure S1D-E). On average, the leg musculature was dim during these periods (Figure S1E), whereas in flexors muscle fibers, which were more superficial, fluorescence could increase more than six-fold during flexion events. GCaMP off kinetics were slow relative to the extension movements (Figure 1E and S1E), and with widefield imaging, diffuse emission from the bright and slowly fading flexors may have obscured small increases in extensor fluorescence. We still found it curious that contractions of extensors did not produce brighter events. We speculate that calcium influx and contractile forces may be larger in flexor muscles than extensors because flies use flexion of the forelimb tibia to support their weight, to hold onto the substrate, and to pull their body during walking, whereas extensors generally lift up unloaded limbs when swinging them forward.

### Whole-cell patch clamp electrophysiology

To perform whole-cell patch clamp recordings, we first covered the fly in a drop of extracellular saline and dissected a window in the ventral cuticle of the thorax, exposing the VNC. The perineural sheath surrounding was ruptured manually with forceps, near the midline, anterior to the T1 neuromeres. To access the motor neuron cell bodies, we first used a large bore cleaning pipette (∼7-10 μm opening) to remove debris and gently blow cell bodies apart to clear a path from the ruptured hole in the sheath to the targeted motor neuron soma. The saline solution was composed of (in mM) 103 NaCl, 3 KCl, 5 TES, 8 trehalose, 10 glucose, 26 NaHCO3, 1 NaH2PO4, 4 MgCl2, 1.5 CaCl2. Saline pH was adjusted to 7.2 and osmolality was adjusted to 270-275 mOsm. Saline was bubbled with 95% O2 / 5% CO2. The recording chamber was then transferred to the microscope and perfused with saline at a rate of 2-3 mL/min.

Whole-cell patch pipettes were pulled with a P-97 linear puller (Sutter Instruments) from borosilicate glass (OD 1.5 mm, ID 0.86 mm) to have approximately 5 MOhm resistance. Pipettes were then further pressure-polished (Goodman and Lockery, 2000) using a microforge equipped with a 100X inverted objective (ALA Scientific Instruments). The final pipette resistance was approximately 12 MOhms. The polished surface allowed for high seal resistances (> 50 GOhms) to limit the impact of seal conductance on V_rest_ (< 1 mV). We used a Multiclamp 700A amplifier (Molecular Devices) for all recordings. The bridge resistance was balanced before sealing onto a soma. The pipette capacitance was compensated after the seal was made. The internal solution for whole-cell recordings was composed of (in mM) 140 KOH, 140 aspartic acid, 10 HEPES, 2 mM EGTA, 1 KCl, 4 MgATP, 0.5 Na3GTP, 13 neurobiotin, with pH adjusted using KOH to 7.2 and osmolality adjusted to 268 mOsm.

The liquid junction potential for the whole cell recordings was −13 mV (Gouwens and Wilson, 2009). We corrected the membrane voltages reported in the paper by post hoc subtraction of the junction potential.

We purchased TTX from Abcam and Methyllycaconitine citrate from Sigma-Aldrich. To stimulate spontaneous movements, we sometimes bath applied 0.5 mM caffeine (Sigma-Aldrich) during whole cell recordings, which prolonged periods of struggling in response to the epifluorescent light. Figure 5 includes trials recorded in caffeine: 45 of the 90 trials from one fast cell, and 50 of the 100 trials from a second fast cell. Caffeine’s effects have been reported to be due to dopamine receptor agonism (Nall et al., 2016). Caffeine application did not increase activity on the leg electrodes during periods of rest.

### Extracellular recordings from leg motor neurons

We recorded electrical activity in the leg by inserting finely pulled glass electrodes (OD 1.5 mm, ID 0.86 mm) into the femur, through the cuticle, taking care to avoid impaling the muscle. The electrodes were filled with extracellular saline. Extracellular currents were recorded in voltage clamp to improve signal to noise. We confirmed that injecting currents of similar size did not move the leg nor produce additional electrical activity. The currents we recorded likely reflect spikes from the motor neuron axons, commensurate with the short latencies between somatic spikes and the events on the leg electrode (Figure 4). In cases where we observed extracellular leg spikes matched to somatic spikes, we then never observed unmatched somatic spikes, which we may have expected if we were observing muscle action potentials. For simplicity we refer to activity recorded in the leg as the electromyogram (EMG), consistent with the use of the term in other organisms.

The content of EMG signals recorded in the leg depended on the placement of the electrode. To improve our chances of recording spikes from specific neurons, we used femur bristles (macrochaetes) as landmarks to place the electrode near the branched axon terminals. These include four large distal macrochaetes (“bristles 0-3”) and one very proximal, called here the “terminal bristle”. Fast motor neuron spikes were most likely found by impaling the leg between bristle 2 and 3, while large intermediate spikes were often found near the terminal bristle. Even still, the polarity, amplitude, and shape of events from identified neurons could vary substantially. When the electrode was placed near the third macrochaete, we tended to record very large units of >200 pA from the fast motor neuron. Fast neuron units were by far the largest amplitude events in the femur. When the electrode was placed in the proximal part of the femur, near the terminal bristle, the spikes from the fast neuron tended to be smaller but still identifiable. We could not unambiguously detect EMG units associated with Flexor 3 in Figure 1. We ran our spike detection routines (see below) on EMG records only in cases where they could be clearly identified, though we could also identify units as coming from specific neurons when EMG spikes aligned with the spikes in the motor neuron cell body.

### Optogenetic activation of leg motor neurons during electrophysiology

We used optogenetic stimulation during electrophysiology experiments to measure the force production of motor neurons and to measure sensory feedback to motor neurons. We drove the expression of CsChrimson (Klapoetke et al., 2014) with 81A07-Gal4, and the expression of Chrimson88 (Strother et al., 2017) with 22A08-Gal4. CsChrimson expression in 22A08-Gal4 prevented straightening of the wings and caused the front legs to be bent midway through each segment. We drove expression of Chrimson (Klapoetke et al., 2014) in sensory neurons with *iav-*LexA while recording from GFP-labeled motor neurons. During recordings we placed a fiber coupled 105 μm cannula (Thorlabs) next to the ventral window in the cuticle and illuminated the T1 neuropil with a 625-nm LED (Thorlabs). We delivered short flashes of 10 or 20 ms to activate neurons, increasing intensity by increasing the voltage supplied to the LED driver (Thorlabs). We measured the power output of the LED for each voltage we used.

### Spike detection from whole-cell recordings and EMG recordings

To detect spikes in current clamp recordings of membrane potential, we applied the following analysis steps to our records of membrane voltage: 1) filter, 2) identify events with large peaks above a threshold, 3) compute a distance from a template for each event, 4) compute the amplitude of the voltage deflection associated with the filtered event, 5) select spikes by thresholding events based both on the distance to the filtered template (< threshold) and on the amplitude of the spike in the voltage record (> threshold). The parameter space for each of these steps was explored in an interactive spike detection interface which can be found at https://github.com/tony-azevedo/spikeDetection.

1. Raw records of membrane potential digitized at 50 kHz were high-pass filtered with a 3-pole Butterworth filter with a cutoff at 898 Hz, low-pass filtered with a 3-pole Butterworth filter with a 209 Hz cutoff, then differentiated and multiplied by −1. Empirically derived, this procedure resulted in a large positive peak associated with the rapid reversal of the membrane potential at the top of a spike.
2. Events were identified by threshold crossing. Thresholds were in the range of 2-10 *10^-5 mV/s. The threshold was set as low as possible to reject baseline noise. Changes in membrane voltage associated with current injection and large secondary peaks associated with the filtered spike waveform oscillations often passed above threshold.
3. Events rising above threshold were then compared with a template using a dynamic time warping procedure to calculate a distance over a 251 sample window centered on the event. The template was calculated for each cell, and though the amplitude varied with spike amplitude, the template shape was generally similar across cells and genotypes.
4. The voltage fluctuations in the raw voltage trace in the 251 sample window surrounding each detected event (the spikes) were then aligned, averaged and normalized to create an estimated spike waveform. Each voltage fluctuation was then projected onto the spike waveform to calculate an amplitude. Thus, each detected peak in the filtered data had an associated distance and amplitude.
5. Finally, events with small distances from the template (< distance threshold), and with large amplitudes (> amplitude threshold) were identified as spikes. Distance thresholds were in the range of ∼8-12 a.u. while amplitude thresholds were in the range of ∼0.5-0.75 for slow neurons and ∼0.8-2 for intermediate and fast neurons.

Since we were interested in measuring spike latencies and conduction velocities, we calculated the second derivative of the raw voltage trace, smoothed over 5 samples, for each identified spike. We found the peak of the second derivative, using that point as the onset/acceleration of the spike. Finally, we inspected records by eye to reject occasional false positives, such as changes in membrane voltage associated with current injection onset.

For a given cell, we applied the same parameters of the spike detection algorithm to all records, with the exception of large current injections into slow motor neurons. In these cases, while spike frequency clearly increased and the leg moved, the spikes became very small and hard to identify. In such cases we hand tuned the parameters and inspected the identified spikes by eye to arrive at estimated spike rates. False positives and negatives were likely increased in these cases. This can be seen in the large spike rate fluctuations in Figure 4, which we did not see when the spike rate was modulated by sensory feedback instead of current injection. Fast, unloaded movements of the leg (Figure S5C-F), also caused large membrane potential fluctuations that made spike detection in slow neurons difficult, we speculate because of large time-varying synaptic conductances that decreased input resistance. We used the same algorithm to detect spikes from EMG current records in cases where EMG events could be readily identified, though the thresholding parameters differed substantially.

### Immunohistochemistry

Following the recording, we dissected and placed the VNC with the legs and head attached in 4% paraformaldehyde in phosphate-buffered saline (PBS) for 20 min. We then separated the legs and head, retaining the right leg with the filled motor neuron and the VNC. The tissue was then washed in PBST (PBS + Triton, 0.2% w/w), placed in blocking solution (PBST + 5% normal goat serum) for 20 min, and then placed for 24 hrs in blocking solution containing a primary antibody for neuropil counterstain (1:50 mouse anti-Bruchpilot, Developmental Studies Hybridoma Bank, nc82 s). After again washing in PBST, the tissue was then placed in blocking solution containing secondary antibodies for 24 hr (streptavidin AlexaFluor conjugate from Invitrogen, 1:250 goat anti-mouse AlexaFluor conjugate from Invitrogen). To label muscles, the leg was fixed, washed, and incubated in blocking solution, like the nervous system. Then it was placed in PBST containing 0.01% sodium azide (Thermo Fisher) and 1 unit of phalloidin (Thermo Fisher) and allowed to incubate for two weeks at 4°C, with occasional nutation. Following staining, the tissue was mounted in Vectashield (Vector Labs) and imaged on a Zeiss 510 confocal microscope (Zeiss). Cells were traced in FIJI (Rueden et al., 2017; Schindelin et al., 2012), using the Simple Neurite Tracing plug-in (Longair et al., 2011). Images in Figure 2 show the results of the filling function in the FIJI plug-in. In some cases, the neuropil counterstain (anti-Bruchpilot) was omitted and the native autofluorescence of the tissue (along with nonspecific binding of streptavidin and GFP fluorescence) was used as reference. To quantify morphology, we measured the soma diameter, the width of the neurite entering the neuropil, and the width of the axon, as close to the exit of the neuropil as possible. We made two measurements for each image and location and averaged the values (Table S1).

### Fly preparation for walking experiments

Fly wings were clipped under cold anesthesia (<4 mins) 24 hours before walking experiments. the fly’s dorsal thorax was attached to a tungsten wire (0.1 mm diameter) with UV-curing glue (KOA 300, KEMXERT). Tethered flies were then given only water for 2-5 hours prior to being placed on the treadmill. In some experiments, the tethered flies were then decapitated under cold anesthesia and allowed to recover for 5-20 minutes prior to the experiment.

Intact tethered flies were positioned on a hand-milled foam treadmill ball (density: 7.3 mg/mm^3^, diameter: 9.46 mm) that was suspended on a stream of air (5 l/min) and freely rotated under the fly’s movement (Figure 7D). The ball and fly were illuminated by three IR lights to improve motion tracking. In experiments on headless flies, we removed the spherical treadmill, leaving the flies suspended in air. For all trials, the temperature in the chamber was maintained between 26-28 °C with a relative humidity of 58-65%.

### Tethered behavior assay

We coaxed flies to walk on the ball by displaying visual stimuli on a semi-circular green LED display (Reiser and Dickinson, 2008). To elicit forward walking, we displayed a single dark bar (width 30°) on a light background, and sinusoidally oscillated the bar at 2.7 Hz across 48.75° about the center of the fly’s visual field. During periods between trials, the LED panels displayed a fixed dark stripe (30°) on a bright background in front of the tethered fly.

To characterize the role of the motor neurons in behaving tethered flies, we optogenetically activated or silenced genetically targeted motor neurons. A green laser (532 nm, CST DPSS laser, Besram Technology, Inc), pulsed at 1200 Hz with a 66% duty cycle, passed through a converging lens and a pinhole (50 µm diameter) with a resulting power of 87 mW/mm^2^ at the target. It was aimed at the fly’s left thoracic coxa-body wall joint, thus targeting the motor neuron axons and the T1 neuromere below the cuticle. Experiments using a driver line labeling all motor neurons (OK371-Gal4) indicated that optogenetic stimulation primarily effected neurons innervating the left prothoracic leg, though we cannot rule out effects on other VNC neurons (Movie S8).

Each trial was four seconds long. We presented walking flies with the visual stimulus, the flies reached a steady running speed at ∼1.5 sec (**Figure 7E**), and the laser stimulus began at 2 seconds. We omitted the laser in some trials (0 ms), or the laser stimulus was either 90 ms or 720 ms in duration, interleaved in random order. Trials were separated by a 25 second period during which video data was written to disk and the LED panels displayed a fixed, stationary stripe.

### Quantification of fly behavior

We used Fictrac (Moore et al., 2014) to calculate fly walking trajectories (position, speed, and rotational velocity) from live video of the spherical treadmill’s rotation (Point Grey Firefly camera, imaging at 30 Hz). Trajectories were then converted from pixels to mm using the spherical treadmill’s diameter of 9.46 mm. Detailed fly movements and kinematics were captured from six simultaneously triggered cameras (Basler acA800-510µm, imaging at 300 Hz) that were spatial distributed around the fly’s body. Digital and analog data signals were collected with a DAQ (PCIe-6321, National Instruments) sampling at 10 kHz and recorded with custom MATLAB scripts.

We manually tracked the position of the left femur and tibia in headless suspended flies from the high-speed video during the 720 ms optogenetic stimulus period using manual annotation in FIJI (Rueden et al., 2017; Schindelin et al., 2012). We then calculated the femur-tibia joint angle from the position measurements in MATLAB.

For each trial on the ball, we scored the fly’s behavior in the 200 ms preceding the optogenetic stimulus as Stationary, Walking/turning, Grooming or Other. Flies that took no steps for the duration of the categorization period were classified as Stationary. Flies that took at least four coordinated steps over the duration of the 200 ms period were classified as Walking/turning, irrespective of any distance traveled. Trials in which the fly switched behaviors, groomed or did not display clear markers for walking/turning during the categorization period were classified as Other/Grooming and excluded from analyses.

For Walking data, we calculated the average forward velocity over time for each stimulus length, for each fly. We computed the percent change in walking speed by averaging walking speed during the stimulation (90 ms and 720 ms) and subsequent 200 ms period for each fly, subtracting the average walking speed at stimulus onset, then dividing by the walking speed at stimulus onset. For Stationary trials, we calculated the percent of trials in which stationary flies reached steady state walking, i.e. sustained walking (>3 mm/s) in the half second following laser stimulation.

We noticed that the laser stimulus caused the empty Gal4 flies (BDP-Gal4) to decrease their walking speed slightly, likely because the flies could see the stimulus. This was particularly noticeable when we used an LED with a large spot size, so we attempted to minimize this behavioral artifact by focusing a laser to a small spot on the fly’s left leg. Even so, the optogenetic stimulus increased the probability that stationary flies would start walking (**Figure 7H**). To control for this effect, we compared the change in speed in motor neuron lines for each stimulus duration, to the change in speed in the control empty Gal4 flies (see statistical analysis below). In no-light trials, walking initiation (**Figure 7H**, black and grey bars) and changes in speed (**Figure 7E**, black traces) did not vary across different genotypes, although baseline walking speed varied slightly across the different lines.

### Statistical analyses

For electrophysiology and calcium imaging results in **Figures 1-6**, no statistical tests were performed *a priori* to decide upon sample sizes, but sample sizes are consistent with conventions in the field. Unless otherwise noted, we used the non-parametric Mann-Whitney-Wilcoxon rank-sum test to compare two populations (e.g. Figure 4) and 2-way ANOVA with Tukey-Kramer corrections for multiple comparisons across three populations (e.g. Figure 3). To compare changes in fluorescence across multiple clusters and extension vs flexion (Figure 1), we used a 2-way ANOVA modeling an interaction between clusters and flexion vs. extension, with Tukey-Kramer corrections for multiple comparisons. To compare cluster ΔF/F of multiple clusters (Figure S1), we used a 2-way ANOVA with Tukey-Kramer corrections for multiple comparisons. All statistical tests were performed with custom code in MATLAB.

For fly walking behavior in **Figure 7**, we used bootstrap simulations with 10,000 random draws to compare both the likelihood of stationary flies to start walking, as well as changes in walking speed (Saravanan et al., 2019). *Stationary* trials were assigned a binary value to indicate that the fly began walking (1) or not (0). For a given stimulus duration and optogenetic condition (Chrimson or gtACR), the binary values for the empty Gal4 control flies and a motor neuron line were combined and then drawn randomly with replacement in proportion to the number of trials for each genotype. As a metric, we measured the difference in the fraction of flies that began walking. The p-value was the fraction of instances in which the randomly drawn distribution produced a value of our metric more extreme then we saw in the data (two-tailed) (**Figure 7H**). For trials in which the fly was already walking at the onset of the laser stimulus (*Walking* trials, **Figure 7E**), we compared the relative change in speed following the stimulus for a given motor neuron line to the empty Gal4 line. We randomly assigned trials to each genotype and calculated the average speed change as above. We used the Benjamini-Hochberg procedure to calculate the false-discovery-rate for either activation or silencing.

## Data and software availability

Data will be made available from the authors website. Acquisition code is available at https://github.com/tony-azevedo/FlySound. Analysis code is available at https://github.com/tony-azevedo/FlyAnalysis.

## Table of Genotypes

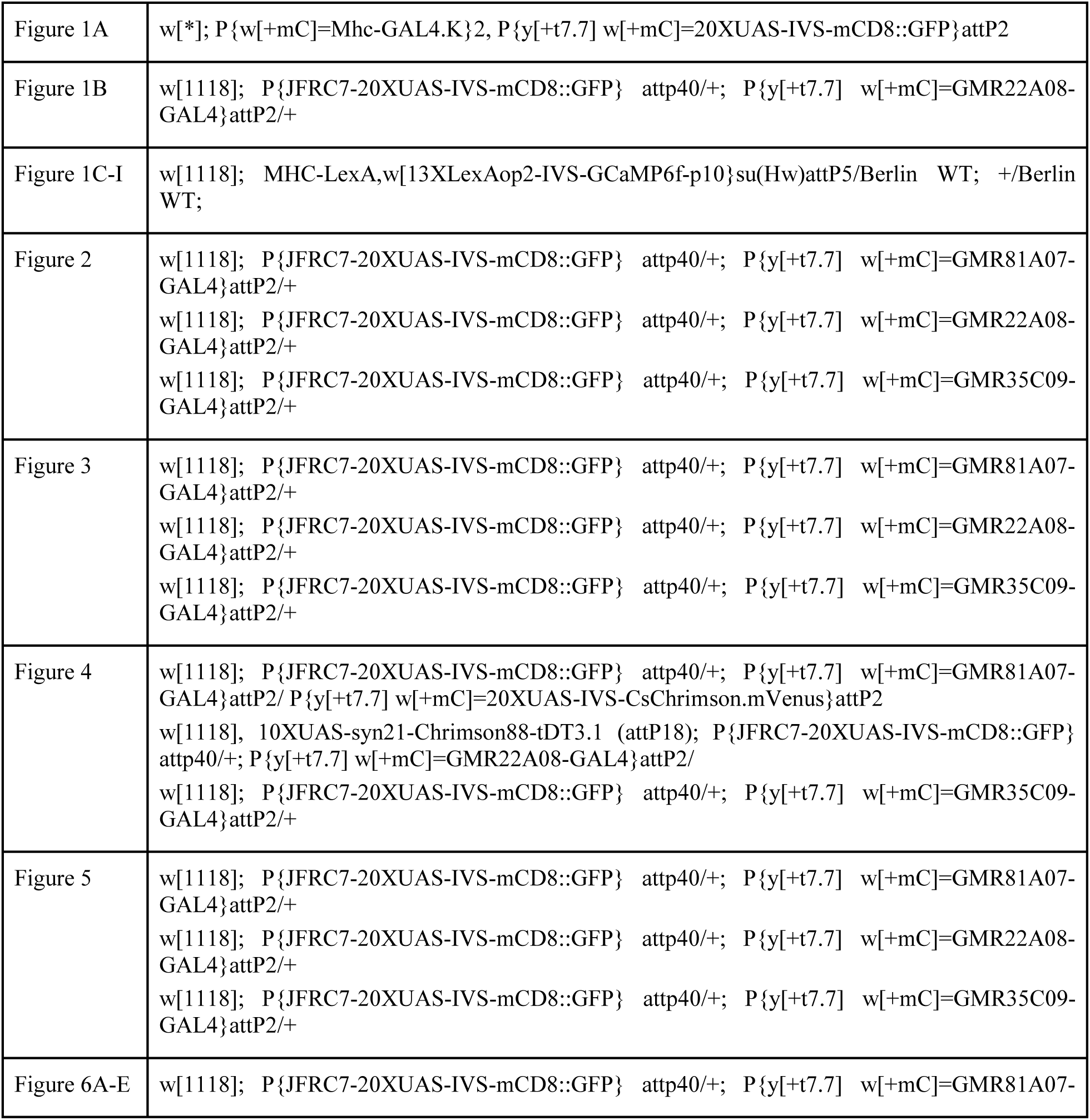

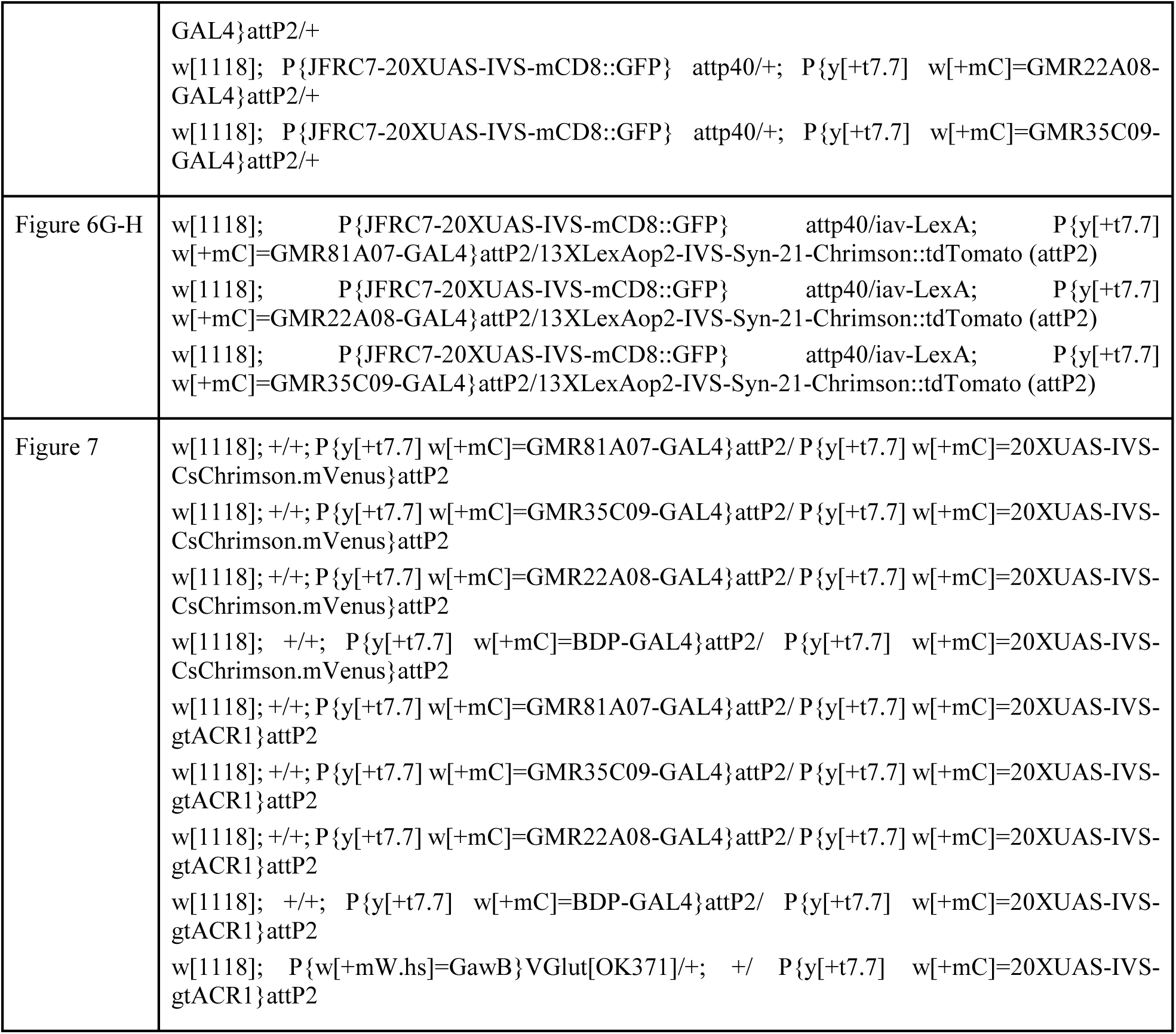

