## Supplemental Figures for "A size principle for leg motor control in *Drosophila*"

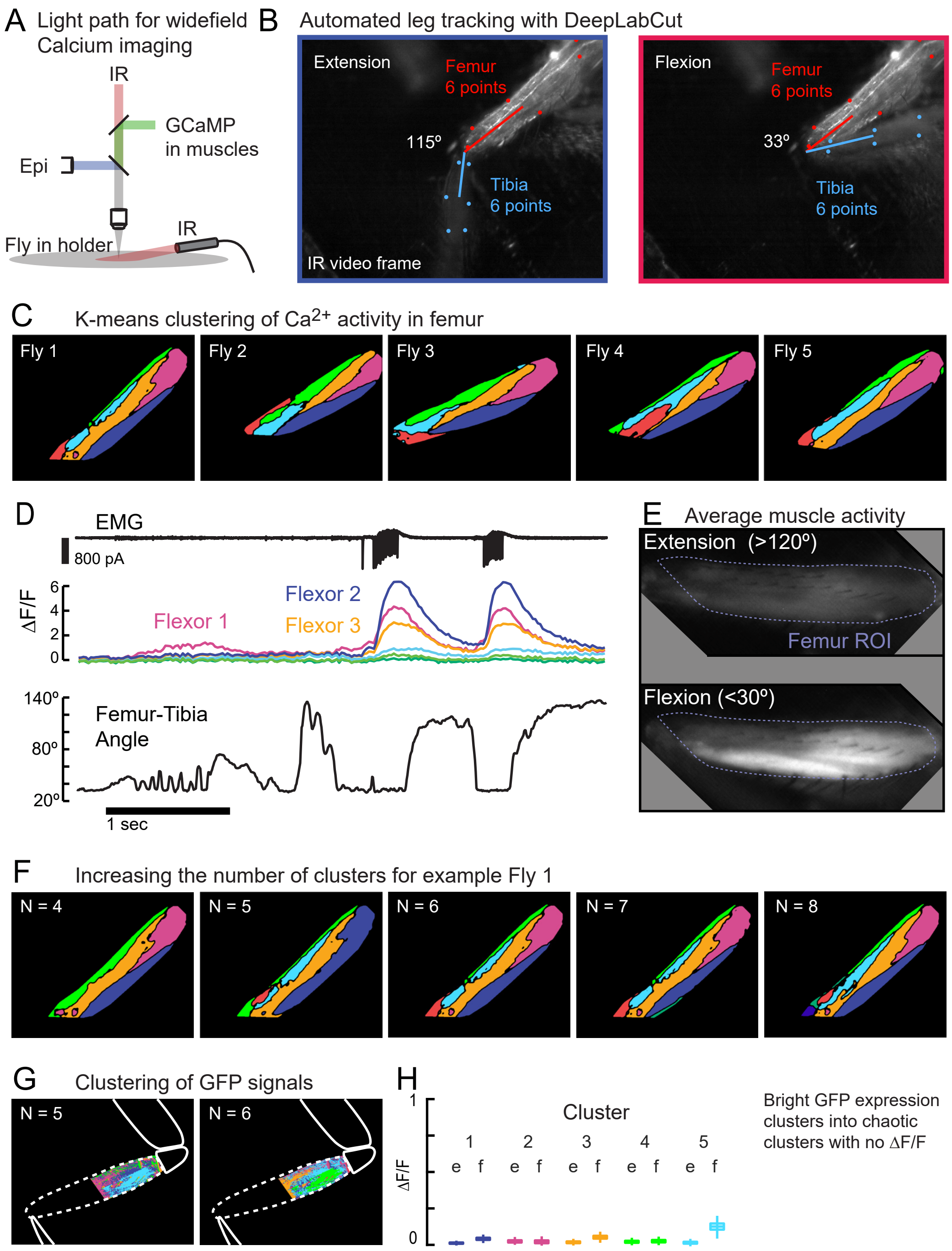


**Figure S1. Wide-field calcium imaging of muscles in the femur.** Related to Figure 1. **A.** Schematic of light path for widefield calcium imaging of femur muscles. Infrared illumination is used to track leg movement. GCaMP emission is reflected by a long pass dichroic mirror to a second camera. **B.** Tracking the femur and tibia with DeepLabCut. A network was trained to detect 6 points on the femur and 6 points on the tibia in the IR illuminated videos of the leg. We calculated the center of the tibia from the 6 detected points. The elliptical arc of the centroid across frames allowed us to estimate the angle of elevation of the tibia relative to the plane of the metal holder (black background). **C.** K-means clustering of GCaMP activity in muscle fibers during unloaded movements of the tibia. Clusters were similar across 5 flies, including the ventral distal cluster 1, the proximal cluster 2, the more dorsal cluster 3, and a thin dorsal cluster 4. Clusters 5 and 6 were more variable in shape and location. **D.** Example epoch from Fly 1 showing the femur-tibia angle and the fluorescence of each cluster. **E.** Averaged fluorescence across all frames in which the tibia was extended (top) vs. flexed (bottom), for Fly 1, normalized to the maximum ΔF/F. Note the lack of signal during extension. **F.** K-means clustering with different numbers of clusters for Fly 1, shown in Figure 1. With fewer than 5 clusters, the proximal cluster tended to be much larger, incorporating much of the region labeled as cluster 3. With more than 6 clusters, the smallest and least modulated clusters tended to divide, not providing any further information. When k=6, pixels in the extension region clustered together, but we did not see large increases in fluorescence with extension (Figure 1). **G.** K-means clustering of GFP expression in muscles during spontaneous movements. Clusters were noisy, interleaved and dispersed. In this case the fly pulled on the probe and the clusters are superimposed over the image of the fly leg. **H.** Changes in fluorescence of clusters calculated from videos of GFP expression during spontaneous movements. GFP expression was bright (Figure 1A), but did not change with movements.

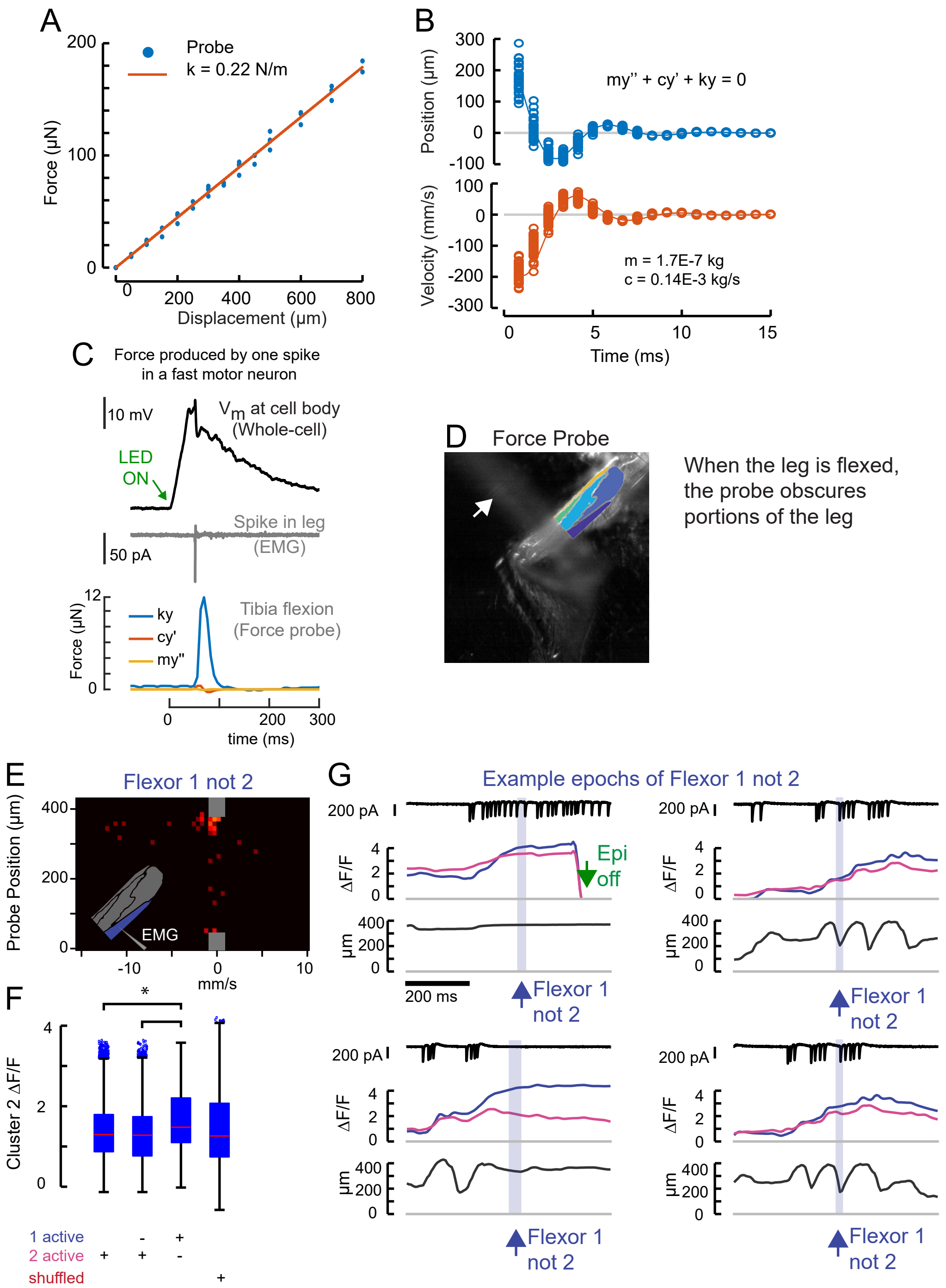


**Figure S2. Calibration of the force probe dynamic properties and muscle activity during probe movements.** Related to Figures 1-4. **A.** The force probe tip was positioned over an analytical balance, and the base of the force probe was displaced. The weight (force) at each displacement is shown in blue, the linear fit is shown in red. The slope of the line is the spring constant, k = 0.2234 N/m. **B.** To measure the effective mass and drag constant of the probe in saline, we flicked the force probe by displacing the tip with a glass hook until it lost contact and snapped back to rest. In blue is the displacement in each frame for 16 different flicks. In orange is the velocity. We used the ode45 solver in Matlab to fit the mass (m = 0.17 mg) and drag coefficient (c = 0.14E-3 kg/s). **C.** Using the dynamical parameters from B, we calculated the portion of the force due to elastic properties of the force probe (blue), drag (red), and inertia (yellow). Here, a single spike evoked optogenetically in a fast motor neuron produced a small movement of the probe (as in Figure 4A, Movie S2). In Figure 4D-F, we calculated force by including drag and inertia, but in other figures we report leg displacement and the approximation of force, assuming that drag and inertia are negligible. **D.** The probe obscured the distal leg during spontaneous movements, so we limited the ROI for clustering to the proximal leg. **E.** The force probe histogram for frames when only the Flexor 1 was active, scaled according to Figure 1H. **F.** Flexor 2 ΔF/F for all frames across N=5 flies in which 1) Flexor 2 is activating (21962 frames); 2) Flexor 2 alone is activating (6686 frames); 3) when Flexor 1 alone is active (1805 frames); 4) random values of cluster 2 ΔF/F (1805 values). The mean Flexor 2 ΔF/F is higher when Flexor 1 alone is active than when Flexor 2 is active (asterisk, p<10-8, 2-way ANOVA, Tukey-Kramer correction for multiple comparisons), indicating that Flexor 2 is also likely contracting even though ΔF/F is not increasing. **G.** Example epochs in which the derivative of Flexor 1 fluorescence, but not Flexor 2 (blue shading) is positive. These epochs make clear that these periods arise when Flexor 2 and 1 are active and contracting together (Flexor 2 ΔF/F high as in panel F), but the derivative of Flexor 2 ΔF/F is either negative or simply not large. The first example occurs near the end of a trial, when the LED was turned off (green arrow).

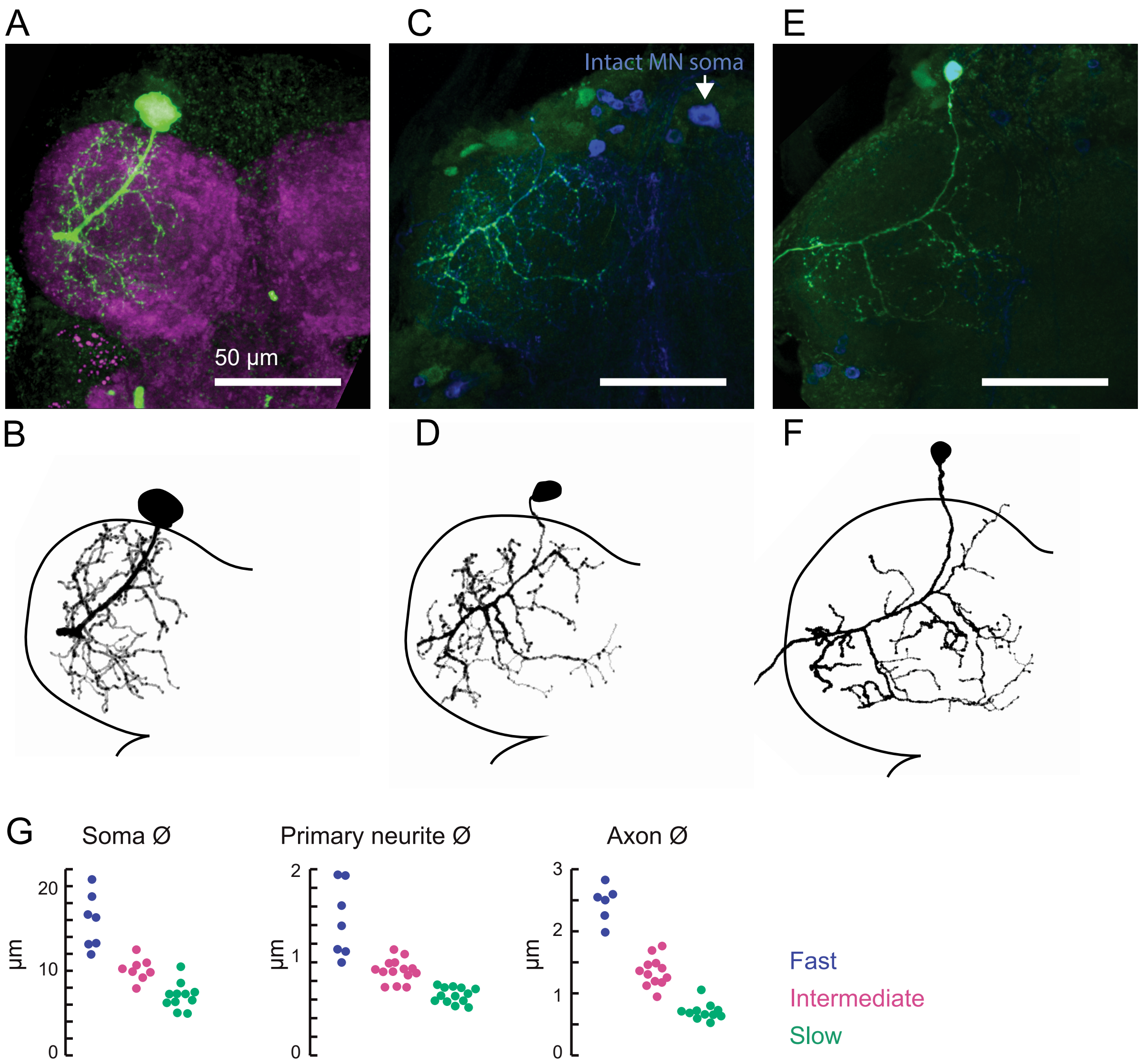


**Figure S3. Central anatomy of flexor motor neurons.** Related to Figure 2. **A.** Maximum intensity projections of fast flexor motor neuron fill (green, neurobiotin). In magenta is the neuropil (nc82). Scale bar is 50 µm. **B.** Traced neuron from A, using the simple neurite tracer plugin in FIJI. Also shown in Figure 2C. **C.** Maximum intensity projections of intermediate flexor motor neuron fill (green, neurobiotin). In blue is GFP expression driven by *R22A08-Gal4*. Note, the cell body of the filled neuron was ripped off, but the motor neuron cell body on the contralateral side remains. **D.** Traced neuron from C, as shown in Figure 2C. **E.** Maximum intensity projections of slow flexor motor neuron fill (green, neurobiotin). In blue is GFP expression driven by *R35C09-Gal4*. **F.** Traced neuron from E, as shown in Figure 2C. **G.** Morphological measurements of identified neurons. Measurements were made of confocal images of filled neurons or GFP-labeled neurons in FIJI. The primary neurite diameter was measured where it crossed the neuropil boundary. The axon diameter was measured at the point at which it exited the neuropil, or the at the last visible spot, if the axon/nerve was torn during dissection of the leg. All groups are different ($p<0.01$, 2-way ANOVA, Tukey-Kramer correction for multiple comparisons).

Table S1: Measurements of motor neuron morphology. Related to Figure 2.

| | *Soma diameter (µm)* | *Primary neurite (µm)* | *Axon (µm)* |
|---|---|---|---|
| *Slow Motor Neurons* | 7.08 ± 2.29 s.d. (10) | 0.65 ± 0.10 s.d. (13) | 0.71 ± 0.14 s.d. (11) |
| *Intermediate MNs* | 10.16± 2.97 s.d. (8) | 0.91 ± 0.14 s.d. (14) | 1.35 ± 0.26 s.d. (12) |
| *Fast Motor Neurons* | 15.84± 4.41 s.d. (7) | 1.45 ± 0.4 s.d. (7) | 2.45 ± 0.31 s.d. (6) |

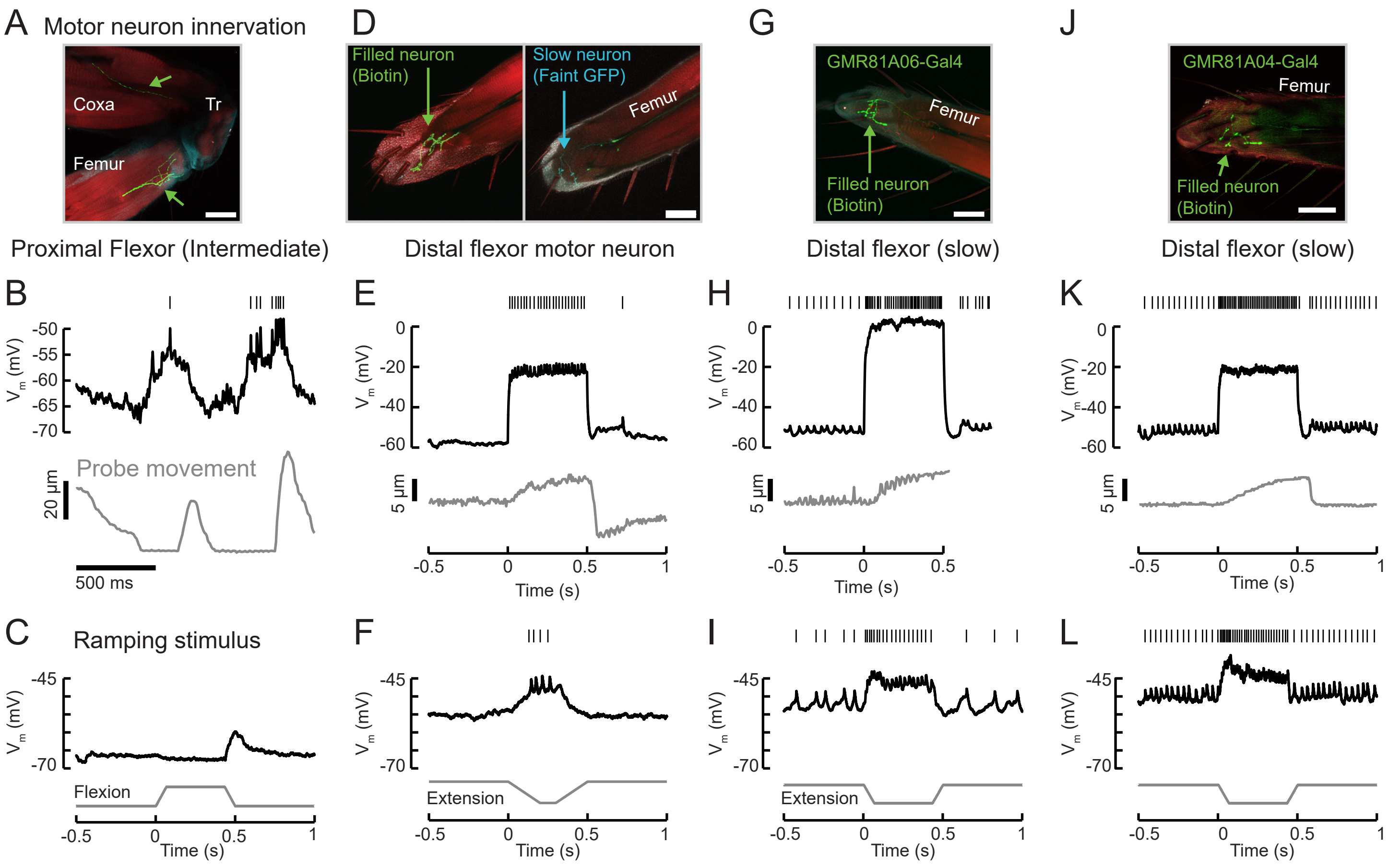


**Figure S4: Example recordings from other tibia flexor neurons.** Related to Figures 4-6. A. Confocal image of an intermediate flexor motor neuron axon (Neurobiotin fill is shown in green; red is phalloidin) from *UAS-GFP;R81A06-Gal4*. The filled neuron targets the proximal femur. Scale bar is 50 µm. The axon appears to target different fibers than the intermediate neuron in *R22A08-Gal4*. **B.** Membrane potential, spikes, and probe movement during spontaneous movements. Membrane potential reflects and slightly precedes flexion events, and spikes occur at low forces. **C.** Single trial example of sensory feedback. The neuron rests at -67 mV. Passive flexion of the leg (60 µm = 8°) hyperpolarizes the intermediate neuron in *R81A06-Gal4*. Extension causes an 8 mV EPSP. Compare to Figure 6. **D.** Confocal image of a flexor motor neuron axon in the leg, neurobiotin fill is shown in green; red is phalloidin. The recording was from an unidentified, non-GFP-expressing neuron in a *UAS-GFP;R35C09-Gal4* fly. The GFP-expressing slow motor neuron is seen in cyan in the right image, at a different z position. The filled neuron targets fibers more proximal than the slow motor neuron. **E.** This neuron rested at -55 mV and did not spike at rest. Current injection could evoke spikes, and moderate spike rates caused steady state tension similar in magnitude to a single intermediate neuron spike (~5 um displacement of the probe). Interestingly, the resulting load on the force probe was sufficient to rapidly force the leg into an extended position. This in turn caused a depolarization and spike in the motor neuron, as the fly returned the probe to the resting position by placing more force on the probe. **F.** Single trial example of sensory feedback. A slow extension of the leg (60 µm = 8°) depolarized the neuron and drove spikes. **G.** Confocal image of a flexor motor neuron axon from *UAS-GFP;R81A06-Gal4*. The filled neuron appears to have a more extensive axon terminal than the slow motor neuron labeled by *R35C09-Gal4*. **H.** This neuron rested around -53 mV, with a spontaneous spike rate of 12 Hz. Large increases in spike rate increased the force on the probe. The recording ended prematurely and we did not capture the probe returning to rest, but note the depolarization and small increase in the spike rate following the end of the current injection, similar to the neuron in panels D-F. Noise in probe position is due to poor lighting and is not time locked to spikes. **I.** Single trial example of sensory feedback. Extension of the leg (60 µm = 8°) of the probe depolarized the neuron and drove spikes. **J.** Confocal image of a flexor motor neuron axon from *UAS-GFP;R81A04-Gal4*. **K.** This neuron rested around -51 mV, with a spontaneous spike rate of 28 Hz, an input resistance of 486 M Ω, and could increase flexion force more than the slow neuron labeled by *R35C09-Gal4*. L. Single trial example of sensory feedback. Extension of the leg (60 µm = 8°) of the probe depolarized the neuron and drove spikes.

**Figure S5: Testing the recruitment hierarchy of motor neurons, in paired recordings and as a function of force probe position and velocity.** Related to Figure 5. **A.** The effective spike rate of fast, intermediate and slow motor neurons, over the course of all trials with spontaneous leg movements. Instantaneous spike rates could be much higher. **B.** From the 2D spike-triggered phase histograms and Bayes' rule, we calculated a likelihood function in order to compare spiking regimes in different neurons. If the force probe has a particular position, p, and velocity, q, the likelihood that a motor neuron fired a spike in the preceding 25 ms (right column, linear color scale from 0-1) is given by the number of frames with p,q that follow a spike (middle column, shown in Figure 5) divided by the full distribution of p,q (left column). The centroids of the full histograms (left column) are shown in Figure 5. **C.** Comparison of activity in recordings of pairs of motor neurons: the number of EMG spikes preceded by a spike in the whole-cell recording, i.e. in a neuron lower down the recruitment hierarchy. The left hand axis indicates the probability of a spike in each millisecond bin for 30 ms before the EMG spike. The right hand axis indicates the cumulative probability of observing a spike in the period before the EMG spike. We found that a small number of intermediate neurons were not preceded by slow neuron spikes (N=110/3082 spikes). **D.** An example trial in which the normal recruitment hierarchy was violated, which only occurred when the leg was unloaded (tibia angle, bottom). During rapid shaking of the leg (highlighted region), the intermediate neuron could spike before the slow motor neuron and have a higher instantaneous spike rate.

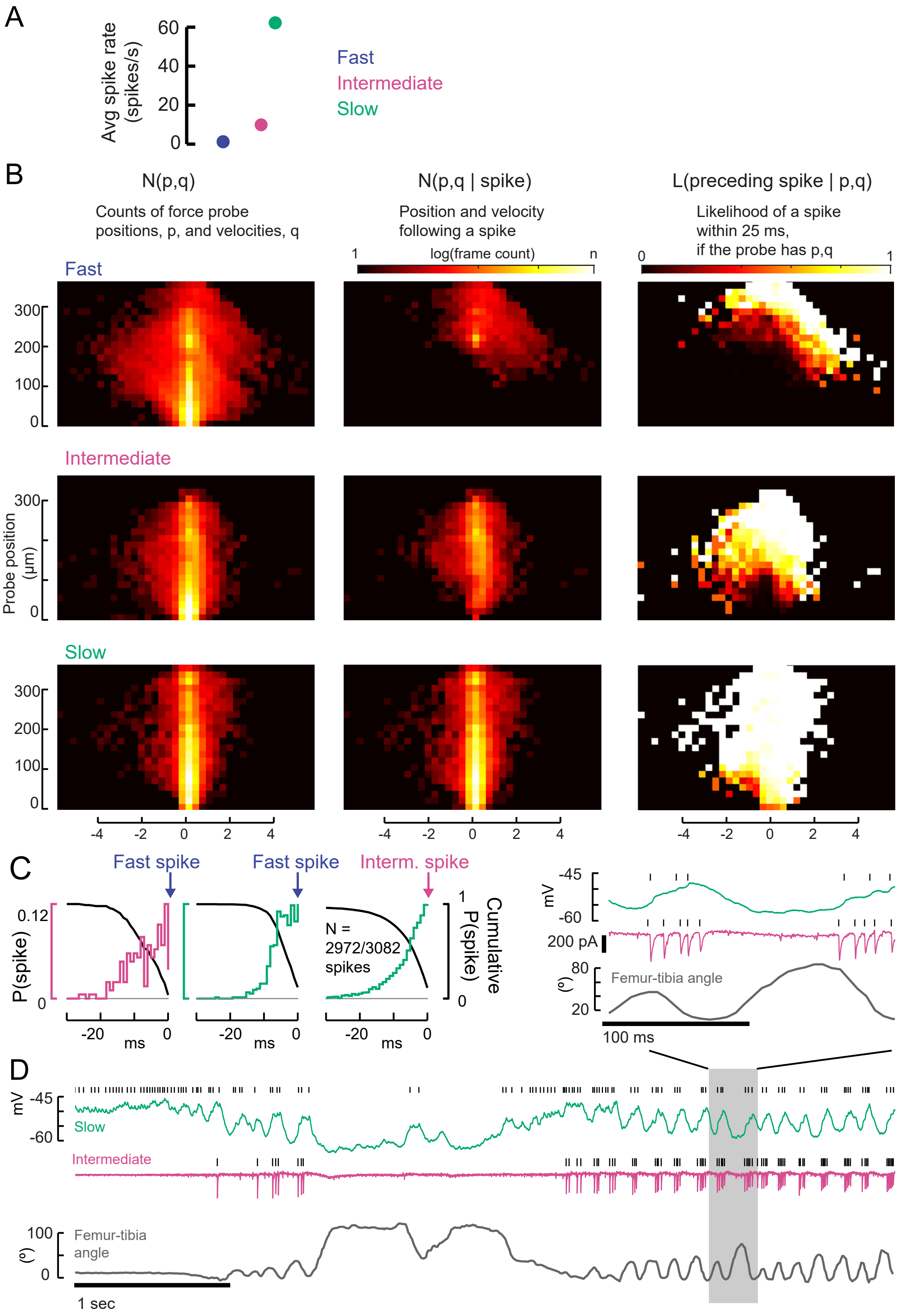

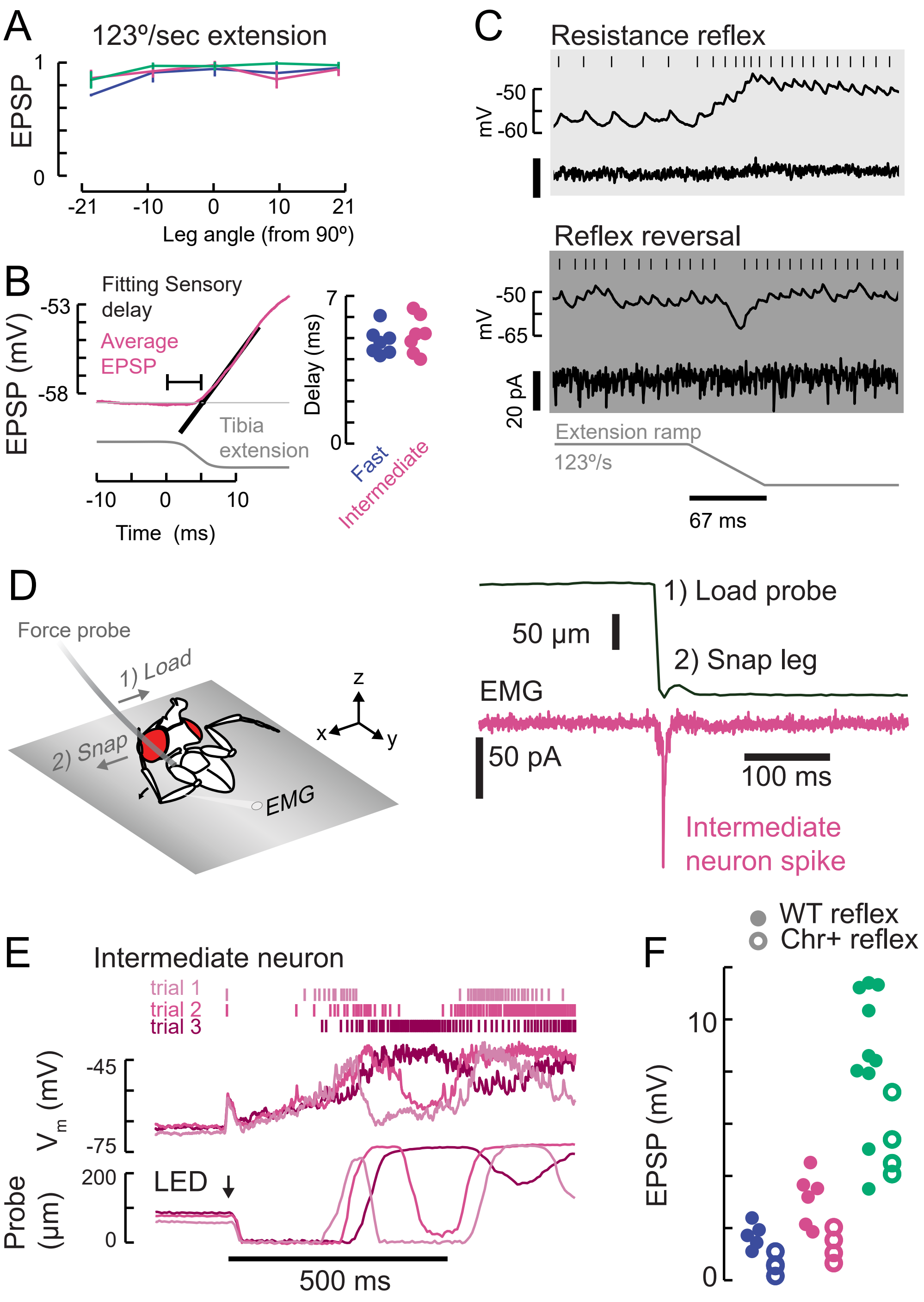


**Figure S6: Properties of sensory feedback to motor neurons.** Related to Figure 6. **A.** The EPSP evoked by fast ramping extension stimuli did not significantly depend on leg angle. **B.** We measured the onset of the sensory evoked EPSP (magenta, intermediate neuron) in the motor neurons by fitting a line (black) to the rising phase of the average response. We measured the time from the start of the command to the piezoelectric actuator for the largest step stimulus (8° extension, shown here as reference). Sensory delays in the fast and intermediate neurons were similar. We did not measure the sensory delays in slow neurons because 1) the spike rate was calculated with an acausal smoothing filter and 2) the average EPSPs were not smooth because of the presence of spikes. **C.** An example of reflex reversal in a slow motor neuron. Top) A normal resistance reflex in which an extension of the leg depolarizes the membrane potential and increases the spike rate. Bottom) When the fly was pulling on the force probe and the EMG activity was increased, the extension stimulus instead hyperpolarized the neuron. **D.** To test faster stimuli than we could deliver with a piezoelectric actuator, we pulled on the probe with a hook until the hook lost contact, and the probe snapped back to rest. We identified intermediate EMG spikes using optogenetic excitation of *R22A08-Gal4* motor neurons. This ballistic stimulus was able to evoke intermediate neuron spikes, but not fast neuron spikes (not shown). **E.** Example responses in an intermediate neuron to a high intensity 10 ms light flash that drove activity in chordotonal neurons expressing Chrimson (*iav-LexA>Chrimson*). The membrane potential and spikes from three trials are shown above (magenta shading) and the simultaneous force probe movement is shown below. Like the flicking stimulus in D, this large stimulus drove intermediate neurons to spike but not fast neurons. The effect of the light stimulus was surprisingly long-lasting, producing membrane potential fluctuations and movements over ~200 ms. **F.** Expression of Chrimson decreases gain of proprioceptive feedback. Peak of the average EPSP in fast (blue), intermediate (magenta), and slow (green) motor neurons in response to a ramping extension stimulus (123 °/sec), in flies that either express Chrimson in FeCO neurons (Chr+ reflex, empty circles) or not (WT reflex, filled circles). Chrimson expression reduces EPSPs, $p<0.01$ for all neurons, Wilcoxon rank-sum test.

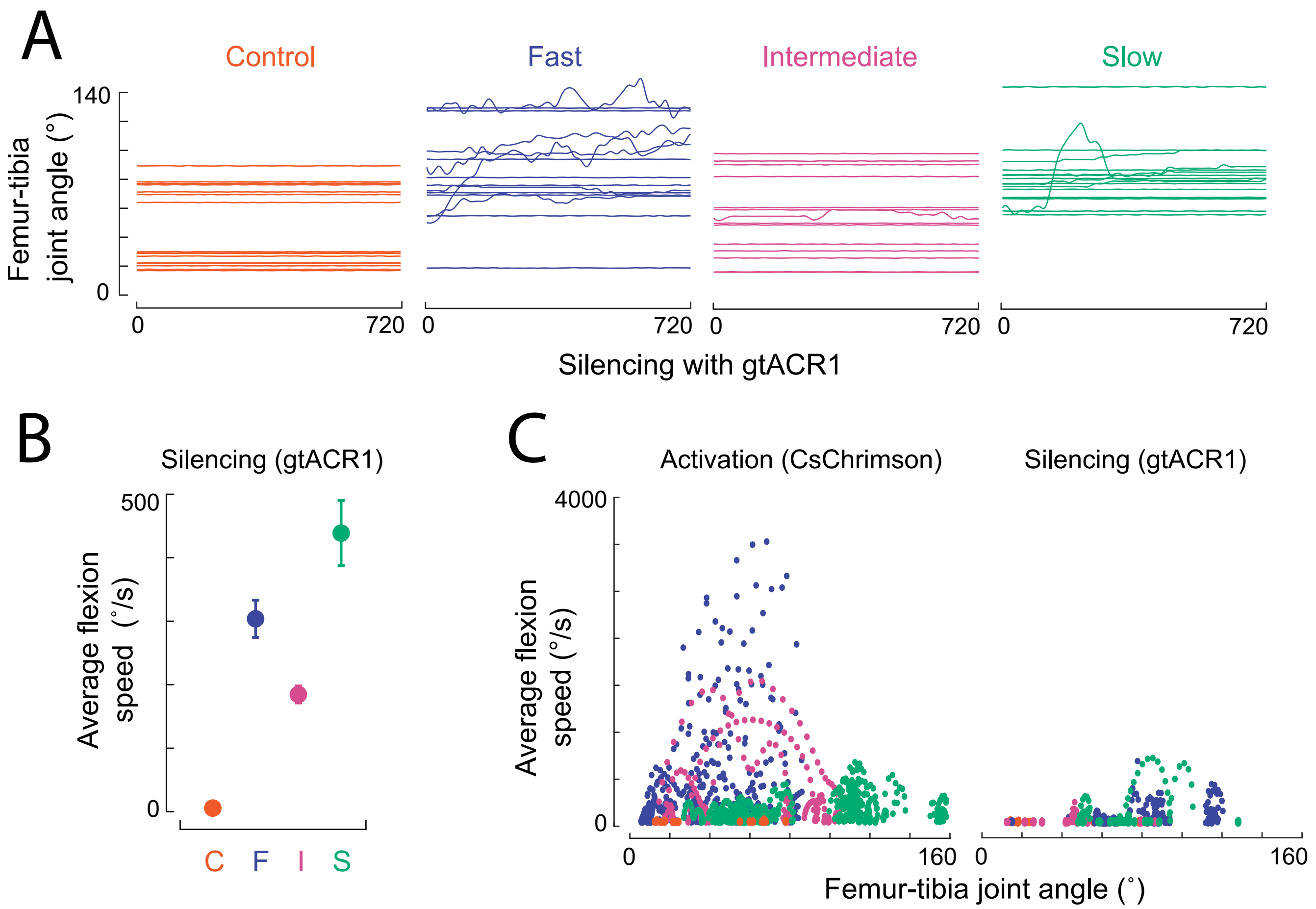


**Figure S7: Effects of tibia motor neuron silencing and activation on leg kinematics.** Related to Figure 7. **A.** Femur-tibia joint angles during silencing (*Gal4>gtACR1*) of the three motor neuron types and a control empty Gal4 driver, in headless, suspended flies. **B.** Average (±sem) flexion speed duringthe initial 300 ms of silencing, on trials where the leg moved. **C.** Scatter plot of instantaneous flexion speed vs. leg angle for each video frame, when activating (left) or silencing (right) the three motor neuron types (and empty gal4 control) in headless suspended flies. Extension would be seen as negative and is not shown. The flexion speeds are largest near 80° for trajectories caused by fast neuron activation (blue points).

**Supplemental Movies available at https://drive.google.com/open?id=1egMTl0ovsiaMKvYU5lZkvsg8D-JwZCof**

**Movie S1. Calcium imaging of femur muscles.** Related to Figure 1, Figure S1 and Figure S2. The video 1) illustrates the arrangement of musculature controlling the fly's tibia; 2) schematizes the position of a restrained fly relative to the force probe fiber, together with wide-field calcium imaging from muscles in the leg and body; and then 3) shows simultaneous videos of the fly pulling on the force with muscle calcium signals in the femur, together with the movement of the probe, an EMG recording from the fast motor neuron, and the time course of cluster ΔF/F. Videos are slowed 3X.

**Movie S2. Motor neuron electrophysiology, force production, and tibia movement.** Related to Figure 4. The video 1) introduces the dendritic morphology and axon projection of the three neurons we studied: fast, intermediate and slow; 2) shows video of individual trials from a fast motor neuron in which one or four spikes are driven with optogenetic stimulation in the VNC, together with probe movement, the whole-cell current clamp recording from the soma, and the EMG record from the leg; 3) shows video of individual trials from an intermediate neuron in which one or four spikes are driven with optogenetic stimulation in the VNC; 4) shows video of a trial from a slow motor neuron in which current injection at the soma depolarizes the neuron and drives ~100 spikes, and a trial in which the slow motor neuron is hyperpolarized, reducing the force on the probe. Videos are slowed 3X.

**Movie S3. Optogenetic activation of motor neurons in headless flies.** Related to Figure 7. The video shows the movements of the flies left front tibia caused by optogenetic activation of the fast, intermediate and slow neurons and a control line (BDP-Gal4). The video shows 1) the tibia flexion during the entire 720 ms laser stimulation period slowed 10X; 2) the first 300 ms of the same event, slowed 40X.

**Movie S4. Optogenetic silencing of motor neurons controlling the front leg.** Related to Figure 7. Motor neurons labeled by *OK371-Gal4* express gtACR1. The video shows the behavior of different flies on different trials while 1) flies were walking during a 90 ms laser stimulus; 2) flies were stationary during a 90 ms laser stimulus; 3) flies were walking during a 720 ms laser stimulus; 4) flies were stationary during a 720 ms laser stimulus. Videos of behavior are slowed 10X.

**Movie S5. Optogenetic activation and silencing of fast motor neurons in behaving flies.** Related to Figure 7. The video shows the behavior of different flies on different trials while 1) flies were walking and the fast motor neuron expressed CsChrimson; 2) flies were stationary, and the fast motor neuron expressed CsChrimson; 4) flies were walking, and the fast motor neuron expressed gtACR1; 3) flies were stationary, and the fast motor neuron expressed gtACR1. Videos of behavior are slowed 10X.

**Movie S6. Optogenetic activation and silencing of intermediate motor neurons in behaving flies.** Related to Figure 7. The video shows the behavior of different flies on different trials while 1) flies were walking and the intermediate neuron expressed CsChrimson; 2) flies were stationary, and the intermediate neuron expressed CsChrimson; 3) flies were walking, and the intermediate neuron expressed gtACR1; 4) flies were stationary, and the intermediate neuron expressed gtACR1. Videos of behavior are slowed 10X.

**Movie S7. Optogenetic activation and silencing of slow motor neurons in behaving flies.** Related to Figure 7. The video shows the behavior of different flies on different trials while 1) flies were walking and the slow motor neuron expressed CsChrimson; 2) flies were stationary, and the slow motor neuron expressed CsChrimson; 3) flies were walking, and the slow motor neuron expressed gtACR1; 4) flies were stationary, and the slow motor neuron expressed gtACR1. Videos of behavior are slowed 10X.

**Movie S8. Optogenetic stimulation of a control line (BDP-Gal4).** Related to Figure 7. The video shows the behavior of different control flies on different trials while 1) UAS-CsChrimson flies were walking; 2) UAS-CsChrimson flies were stationary; 3) UAS- gtACR1 flies were walking; 4) UAS- gtACR1 flies were stationary. Videos of behavior are slowed 10X.
